# Prophage induction states drive structural and synergistic outcomes in marine bacterial biofilms

**DOI:** 10.1101/2025.09.29.679375

**Authors:** Matthew J. Tuttle, Gabi Steinbach, Tapan Goel, Joshua S. Weitz, Alison Buchan

## Abstract

Microbes commonly live in biofilms, dense spatially structured assemblages where temperate phages can influence interactions amongst closely related strains. Yet, how these phage-host interactions impact biofilm architecture and ecological dynamics remains poorly understood, particularly in situations where antagonistic strains coexist in structured communities. In the marine Roseobacteraceae *Sulfitobacter pontiacus*, two genetically similar heteroimmune prophages, (termed A and D) establish lysogeny in their host. Depending on the host plasmid genotype, prophages are fixed in either high or low induction states, which consequently shape both biofilm structure and competitive outcomes. Using confocal scanning laser microscopy of fluorescently labeled lysogens, we investigated how prophage induction and phage genotype of phage-host coalitions influence monoculture and co-culture (competitive) *S. pontiacus* biofilms grown at a fluid-solid interface. Compared to their low induction counterparts, high-induction strains formed thicker and more voluminous monoculture biofilms, which primarily result from vertical protrusions arising from a basal monolayer. In co-culture experiments – mixing strains with reciprocal prophage genotypes that can mutually kill each other through phage-mediated allelopathy – we observed unexpected coexistence and synergistic biofilm enhancement rather than competitive exclusion. Reactive oxygen species (ROS) levels were significantly higher in co-cultures than monocultures and correlated with local cell density, suggesting that phage-mediated interactions generate hotspots of oxidative stress, which is expected to further promote prophage induction, creating a positive feedback loop that amplifies cell lysis and release of matrix components. These results suggest temperate phages can paradoxically stabilize and promote biofilm growth via mechanisms that combine antagonistic killing with cooperative matrix production, revealing how phage-mediated interactions can shape microbial community architecture in spatially structured environments.

**IMPORTANCE:** Bacterial biofilms are ubiquitous in natural environments, yet understanding of how temperate bacteriophages shape biofilm development and microbial community dynamics remains limited. Here, we demonstrate that prophage induction state simultaneously drives antagonistic killing and cooperative biofilm enhancement in spatially structured microbial communities. Using *Sulfitobacter pontiacus* and its infecting temperate phages as a model system, we reveal that reciprocally antagonistic host-phage pairs, which can kill each other through phage-mediated allelopathy, form more robust biofilms when grown together rather than separately. These findings challenge traditional views of competition in dense microbial systems, highlighting the complex, multi-faceted roles that temperate phages play in microbial communities. Additionally, we propose that oxidative stress creates positive feedback loops that amplify prophage induction, promoting coexistence of host-phage pairs rather than driving competitive exclusion. These insights have broad implications for understanding microbial community assembly, maintenance of genetic diversity in biofilms, and evolutionary dynamics of host-phage interactions in structured environments.

## INTRODUCTION

Biofilms are complex, three-dimensional communities of microbial cells encased within a self-produced matrix. Biofilm matrices are composed of diverse microbial polymers, including proteins, exopolysaccharides, and extracellular DNA, and offer community members both protection from external stressors as well as access to captured resources (Flemming et al., 2023). To reap the benefits of life within a biofilm, entrained cells must also manage the inherent stresses imposed by the matrix, including oxidative stress, limited diffusion, and competition with other biofilm members (Flemming et al., 2016). In addition, natural biofilms are overwhelmingly composed of diverse microbial species and strains in distinct physiological states (Stewart and Franklin, 2008; Stubbendieck and Straight, 2016). The population dynamics in these dense multispecies communities are determined by interaction networks amongst different members as they engage in both cooperative and competitive interactions for space, resources, and survival (Dorosky et al., 2017; Rostami et al., 2022; Teschler et al., 2022, Cude et al., 2012, Stubbendieck et al., 2016). Consequently, the spatial structure of biofilms and the arrangement of genotypes can influence the type and frequency of interactions (Nadell et al., 2016). For instance, in biofilms with spatially segregated genetic lineages, greater cooperation is expected among proximate genotypic clones; meanwhile, in biofilms with greater spatial mixing, antagonistic interactions may be expected at greater frequency. The physiochemical nature of the matrix and proximity of embedded cells increases the frequency of deadly microbial interactions when different genotypes are present (Abe et al., 2020).

Beyond cell-cell interactions, temperate bacteriophages can also function as bacterial weapons against their closest relatives through a process termed phage-mediated allelopathy (e.g., Basso et al., 2020; Davies et al., 2016). When two heteroimmune lysogenic strains are grown together in co-culture, they may attack one another through the production of competitor-specific phage, which can either initiate a lytic or lysogenic infection in the susceptible competitor (Basso et al., 2020). This form of intraspecific interaction is poorly understood, but potentially pervasive (Harrison and Brockhurst, 2017). As intraspecific interactions appear to dictate eco-evolutionary dynamics in bacterial communities (Goyal et al., 2022), it is critical to understand the mechanisms that drive strain level competition. Phage-host interactions within biofilms are particularly complex, as they have been demonstrated to have positive and negative symbiotic effects (Hansen et al., 2019; Pires et al., 2021). Promotion of biofilm formation by induction of temperate phages has been seen among several phage-host pairs, and often attributed to release of biofilm matrix components, notably extracellular DNA (eDNA) (Carrolo et al., 2010; Gödeke et al., 2011; Nanda et al., 2015; Turnbull et al., 2016). Conversely, lysogeny has also been shown to be detrimental to biofilm formation (Gillis et al., 2014; Liu et al., 2015; Zegans et al., 2009) and induction can be used as a mechanism for biofilm dissemination (Rossmann et al., 2015).

A member of the ecologically important *Roseobacteraceae* family, *Sulfitobacter pontiacus*, is emerging as an environmentally relevant and tractable system for unraveling the complex interplay of host-phage interactions and understanding their biological significance (e.g., Ankrah et al., 2014; Basso et al., 2022; Tuttle et al., 2022). Prior studies using lysogenized strains that are locked in either a high or low prophage induction state showed that induction state influences biofilm biomass and structure (Basso et al., 2020). Moreover, phage-mediated allelopathy, utilizing two genetically similar temperate phages that can independently lysogenize their shared host, drives intra-strain competition, revealing divergent influences on the competitiveness of this bacterium in distinct environmental niches. The outcome of the competition, i.e., which host-phage pair outcompetes the other, differed between planktonic and surface-attached growth (Basso et al., 2020). This asymmetric competition suggests that coexistence of both host-phage pairs could be feasible in heterogeneous environments.

Here we set out to evaluate mechanisms of phage-mediated bacterial strain coexistence building on our foundational work, which was limited to bulk community analyses and unable to differentiate free phage from prophage, *i.e*., lysogenized hosts (Basso et al., 2020). Consequently, strain-specific interactions, notably spatial organization, could not be deduced. To better assess how these phage-host pairs influence conspecific interactions in the context of a biofilm at spatial resolution, here we use confocal scanning laser microscopy (CSLM) of constitutively fluorescent strains to examine biofilms of single and mixtures of *S. pontiacus* strains that exhibit one of two induction states (i.e., high or low). We first analyzed the global and local spatial architecture of monoculture biofilms of lysogenic strains. We then compared the results to the global architecture of co-culture competitive biofilms and the spatial representation of each strain within co-cultures. Finally, we characterized the biofilm matrices. Our results show that phage-host coalitions alter biofilm structure and dictate population outcomes, with reactive oxygen species playing a potentially pivotal role.

## RESULTS

### Study system

Prophage induction and subsequent phage-mediated killing of adjacent cells is expected to play a critical role in spatially explicit interaction dynamics. Here, we utilize a previously described roseobacter model system in which two genetically similar temperate phages, termed here “A” and “D”, influence monoculture competition dynamics in their shared host, *S. pontiacus*, in a growth modality-dependent fashion (Basso et al., 2020). *S. pontiacus* is lysogenized by either prophage genotype, with the resulting lysogens termed CB-A and CB-D, to distinguish between the chromosomally encoded prophage genotype, A and D, respectively. We have previously demonstrated plasmid-dependent stabilization of either prophage in the lysogenic state (Tuttle et al., 2022). More specifically, we showed a robust link between plasmid content and prophage induction. Strains possessing all four native plasmids have undetectable phage in cell-free lysates, referred to here as “low” induction state and are termed CB-D(lo) and CB-A(lo). Strains lacking two plasmids (pSpoCB-2 and pSpoCB-4) represent a “high” induction state as they produce considerable (10^5^ PFU/mL) phage titers, subsequently termed CB-D(hi) and CB-A(hi). To systematically derive the impact of phage-mediated monoculture competition in the co-culture biofilms, we used a combination of lysogen genotype and induction state to create a set of four distinct strains (Fig 1A) where lytic infections occur on hosts with reciprocal prophage genotypes and different plasmid contents.

**Figure 1.**
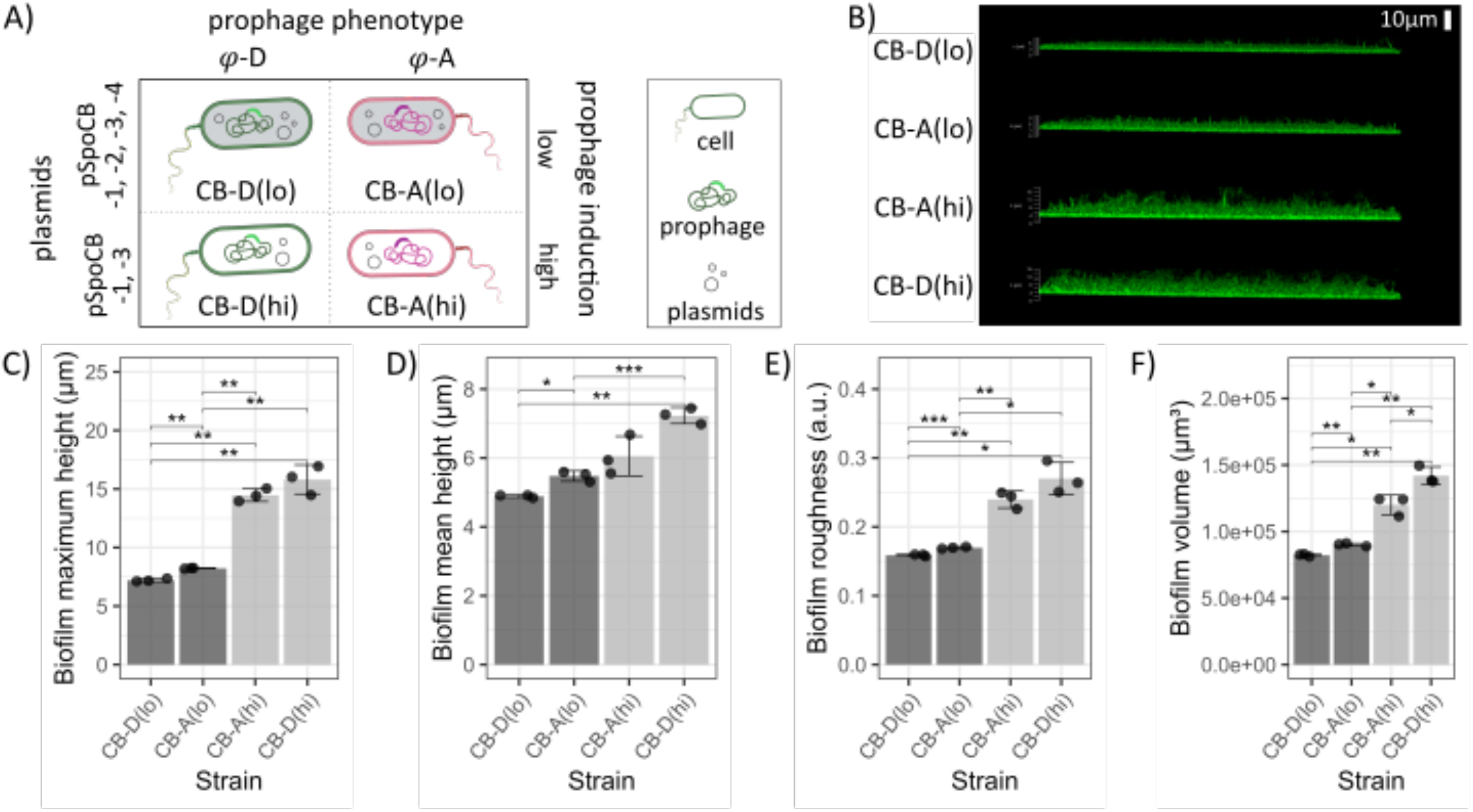
Strains lacking plasmids exhibit higher rates of prophage induction and form thicker, rougher, and more voluminous biofilms in monoculture. **(A)** The four lysogen genotypes in this study differ by prophage genotype (D or A) and number of plasmids (two or four), where number of plasmids impacts induction state (low or high). **(B)** Side views of representative three-dimensional CSLM micrographs of all four, GFP-labeled strains grown for 24 h as monoculture biofilms. **(C)** Maximum height of monoculture biofilms, defined as the height of z coordinates in the range [1%, 99%]. **(D)** Mean height of monoculture biofilms, defined as the mean height of vertical stacks of objects from cube segmentation of images. **(E)** Roughness coefficient of monoculture biofilms. **(F)** Total volume of monoculture biofilms as measured by volume of GFP fluorescence for images of lateral size 185 μm x 185 μm. For panels **C-F** dots represent replicates from Z-stack images, columns represent the mean of all replicates (n = 3), and error bars depict standard deviation. Student’s t-tests were performed to determine statistically significant differences between means (* p ≤ 0.05; ** p ≤ 0.01; *** p ≤ 0.001; **** p ≤ 0.0001). Data for biofilm properties measured in this experiment are available in Tables S1 and S2.

### Prophage induction state alters global architecture in monoculture biofilms

We examined the global features of monoculture biofilm structure across the four strains in Fig 1A. Strains were genetically manipulated via chromosomal insertions to constitutively express GFP. Side views of biofilms indicate different biofilm structure across the four strains (Fig 1B). A quantitative analysis showed that high induction strains produced biofilms with significantly increased biofilm height (maximum and mean height), volume, and surface roughness relative to their low induction counterparts (Fig 1 C-F). The feature with greatest deviation between the two induction states was maximum height: high induction strain biofilms are approximately twice as tall as low induction strain biofilms (~15 µm vs ~7.5 µm). That neither biofilm mean height nor volume show the same level of difference between the two induction states suggests a variation in spatial organization is driving observed global differences. This hypothesis is supported by the variation in biofilm roughness, a measure of surface height fluctuations (Fig 1E). Collectively, these results demonstrate prophage induction state influences biomass formation and structure within *S. pontiacus* monoculture biofilms.

The choice of fluorophore or specific Tn5-insertion sites had no significant influence on biofilm properties. The same trends in biofilm maximum height, mean height, roughness, and volume were observed in a repeated experiment using a separate set of strains expressing mCherry rather than GFP (Fig S1 and Table S1). The comparison of mean values across three replicates indicated small significant differences between GFP- and mCherry-fluorescent strains of the same prophage and induction state genotype (Table S2).

### Monoculture biofilms form basal monolayers from which extensive protrusions emerge in high induction state strains

To examine local spatial structure within high and low induction state biofilms, GFP-expressing CB-D(lo) and CB-D(hi) strain biofilms were considered and resolved at the level of individual cells within the biofilm matrix. Motivated by the contrasting variation in maximum and mean height between low and high inducers (Figs 1C, D), we sought to investigate the height-dependent architecture of the CB-D monoculture biofilms. Vertical cross sections (Fig 2A) reveal that for both low and high inducer strains the bottom of the biofilm consists of a monolayer of cells across the glass surface on which they were attached. *S. pontiacus*, like many other roseobacters, typically appear as dumbbell-like or “matreshka”-shaped cells (Sorokin, 1995). Here, we observe that the surface-attached cells exhibit polar attachment, *i.e*., attach with their cell head to the surface and extend with their long axis into the liquid medium.

**Figure 2.**
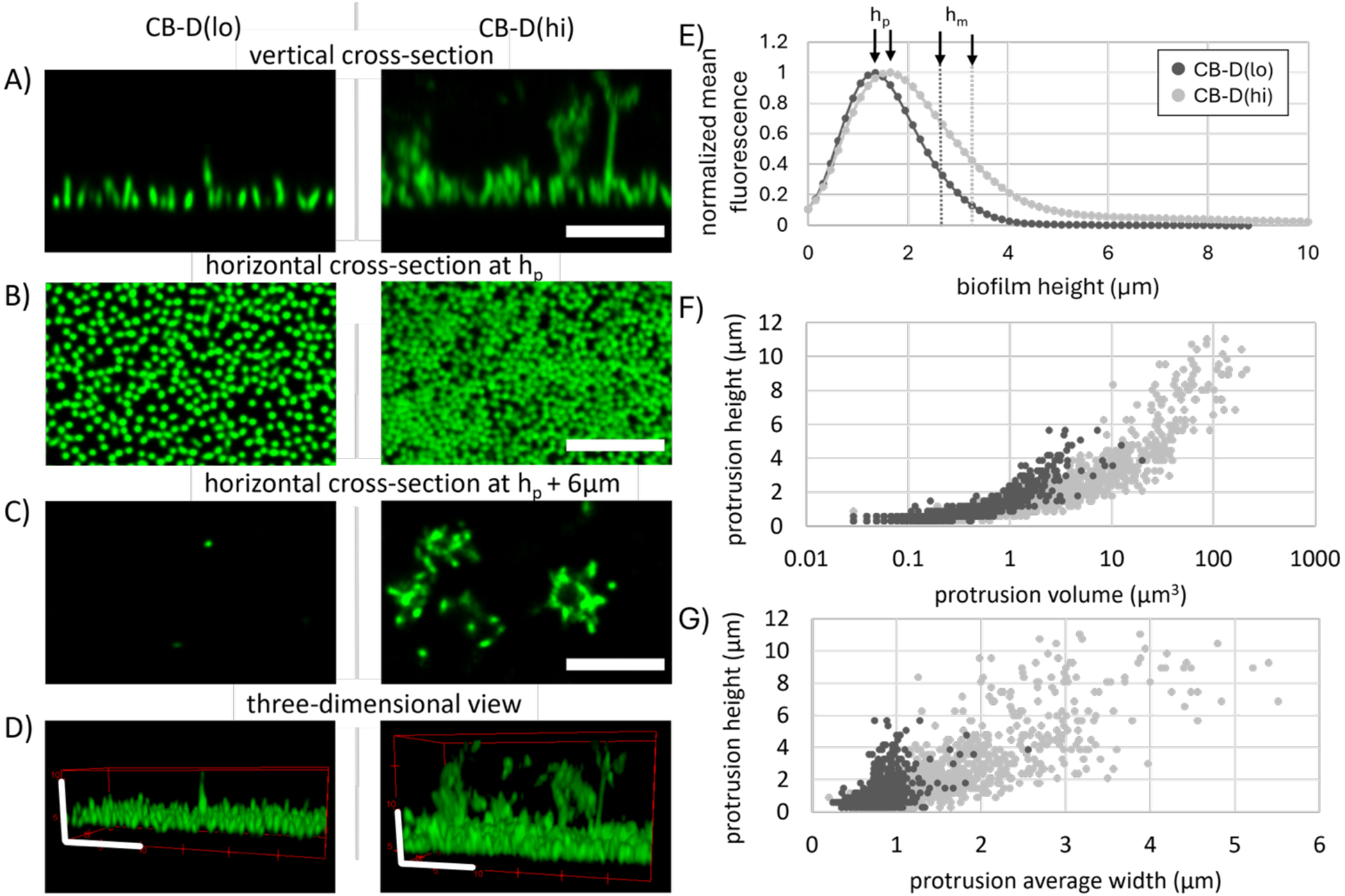
Spatial structure of CB-D(lo) and CB-D(hi) biofilms exhibits a cell monolayer with protrusions of differing size. Representative confocal microscopy images of 24 hour CB-D(lo) and CB-D(hi) biofilms are shown as cross-sections from **(A)** vertical cuts, **(B)** horizontal cuts through the monolayer (at h_p_), **(C)** cuts 6 μm above h_p_, and **(D)** three-dimensional views. White scale bars in panels A-C and D (different lateral and vertical resolution) correspond to 10 μm. **(E)** Normalized distribution of fluorescence intensity as a function of biofilm height averaged over three replicate samples for both CB-D strains. Protrusions above the cellular monolayer, h_m_, are characterized by protrusion height over **(F)** protrusion volume and **(G)** protrusion average width.

Horizontal cross-sections through the monolayer base show that cells are densely packed for both strains, with higher packing in the high-inducer strain biofilm (Fig 2B). Above the monolayer, CB-D(hi) generates pronounced protrusions - extended, but clearly separate three-dimensional clusters of cells - that extend away from the monolayer base into the overlaying aqueous medium (Figs 2A, 2C). In contrast, CB-D(lo) biofilms contain much smaller protrusions, on the order of a single cell width (Fig 2C) and a few cell lengths in height (Figs 2C, 2D). Analyses of strains with the alternate “A” prophage genotype (*i.e*., CB-A) demonstrate the same architectural features and trends between low and high induction biofilms (see supplemental Fig S2). Accordingly, the induction state (i.e., production of phage particles and lysis of producing cells) appears to drive observed differences in biofilm development.

Next, we quantified the height-dependent distribution of cellular biomass by measuring the mean fluorescence intensity as a function of biofilm height (i.e., the distance to the glass surface where cells have attached). The fluorescence intensity curves, normalized by maximum intensity, exhibit a peak at biofilm height h_p_ that is similar for both strains CB-D(lo) and CB-D(hi) (Fig 2E). A notable difference appears at the right side of the curve; the long tail that appears significantly broader for the CB-D(hi) strain. Analyses of strains with the alternate “A” prophage genotype (i.e., CB-A) exhibit the same shapes and trend of intensity curves for low and high induction biofilms (Fig S3). To quantify the two distinct structural elements, i.e., monolayer base and protrusions, we split the curves into two sections: the bottom section around the peak with an upper limit at biofilm height h_m_=2* h_p_, and the top section capturing the tail of the intensity curve at heights h>2* h_p._ The width of the peak, h_m_, is comparable for both strains: on the order of one cell length (differing by < 0.6 µm on average, Table S3), correlating with the dense monolayer of cells with polar attachment for both strains observed in the CSLM images. We further find that the fluorescence intensity at the peak, I(h_p_), varies across strains (Table S3), following the same trend as the extracted biofilm volumes for the strains (Fig 1). The variation in peak fluorescence intensity between CB-D(hi) and CB-D(lo) matches the visual difference in packing density (Fig 2B). Finally, we assessed biomass distribution between the monolayer base and the protrusions. To do so, the integrated fluorescence intensity of the tail (integrated fluorescence intensity for biofilm heights h>h_m_) was divided by the integrated fluorescence intensity of the monolayer section (for heights h<h_m_), defined as the tail-to-monolayer intensity ratio Φ_tm_ (Table S3). We obtained larger Φ_tm_ values in high-induction biofilms as compared to their low inducer counterparts, indicating enhanced relative biomass content in the protrusions.

To compare the architecture of the protrusions in CB-D strains, we extracted and compared their shape characteristics, i.e., volume, height and average diameter (Figs 2F, G). This analysis uses the sub-stack of Z-plane confocal images that contain only the protrusions, above height h_m_. Volumes of protrusions in CB-D(lo) biofilms are about the same as, or less than, the volume of a few single cells, indicating that only a few individual cells grow (or attach to) the biofilm beyond the surface monolayer. Accordingly, the diameter and height of CB-D(lo) protrusions are approximately one cell width by up to three cell lengths, respectively (Fig 2G). This suggests that as individual cells divide along their transverse plane, they either remain attached to their mother cells or are trapped in the biofilm matrix in an upward extending orientation. In contrast, protrusion volumes in CB-D(hi) biofilms are upwards of two orders of magnitude larger than those of their CB-D(lo) counterpart. These protrusions are several micrometers wide and tall and are predicted to house cell assemblages in the tens to hundreds of cells.

### Co-culture biofilms have more biomass and a different spatial distribution of cells

The analysis of global structure and local architecture in monoculture biofilms was used to predict the consequences of phage-driven competitiveness of strains within co-cultured biofilms when strains possess different prophage genotypes and induction states. In the co-cultured biofilm, released D phages from CB-D cells may infect and kill CB-A cells and vice versa (Basso et al., 2020). Here, biofilms were grown from strains of reciprocal prophage genotypes, using the following pairs: (i) CB-D(lo) + CB-A(lo); (ii) CB-D(lo) + CB-A(hi); and (iii) CB-D(hi) + CB-A(hi) (Fig 3). For each pair, three different inoculation ratios were used to study head-to-head competition (1:1) and invasion-from-rare competition (5:1 and 1:5) dynamics. Irrespective of mixing ratios, all biofilms were seeded with the same absolute concentration of cells. Strains were distinguished by their fluorescent marker: GFP for D strains and mCherry for A strains.

**Figure 3.**
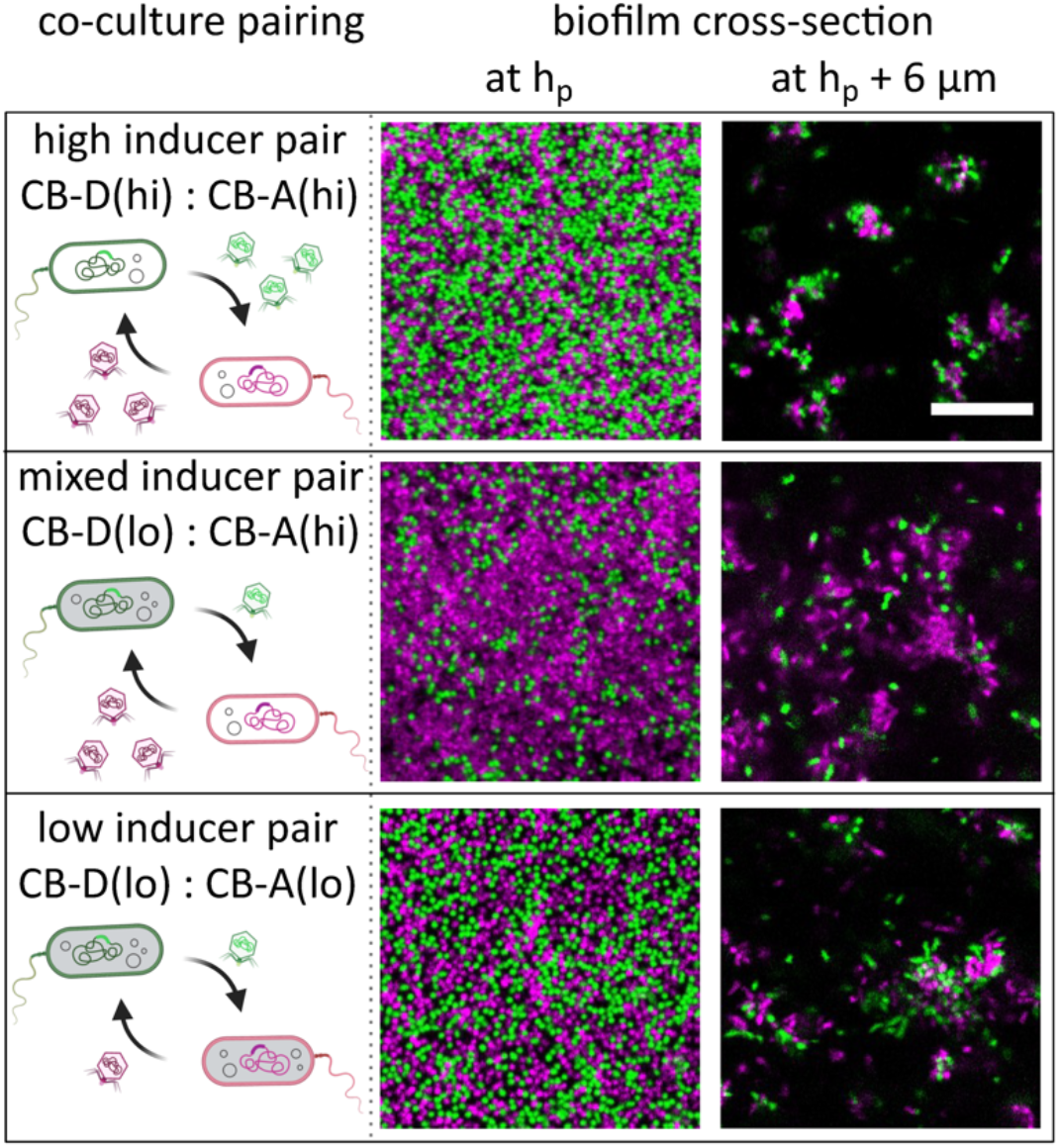
Co-cultured biofilms grown from pairs of strains of varying induction levels and prophage genotypes at 1:1 inoculation ratio indicate synergistic interactions. Co-culture pairing mixes cells from reciprocal phage genotypes that can mutually kill each other through released D and A phages. Horizontal cross-sections of biofilms recorded at the middle of the monolayer base (height h_p_) and 6 μm above demonstrate the biofilm architecture. Scale bar is 15 μm.

To gain insight into the competitive interaction, we recorded cross-sectional confocal images of biofilms at heights near the fluorescence intensity peak, h_p_ (in the middle of the monolayer base), and 6μm above h_p_ for all three co-culture pairs at 1:1 inoculation ratio (Fig 3). We find that in all cases (i) the architecture of a dense monolayer with protrusions atop is retained, and protrusions are very pronounced, (ii) co-cultured strains coexist at 24h, and (iii) cells from the co-cultured strains intermix, showing no clear indication of spatial assortment amongst the strains.

We quantified the height dependent distribution of cells within the co-cultured biofilms. Here, we focus on the outcome for the GFP-labeled CB-D strains in the co-cultured biofilms. For CB-D cells, we extracted the thickness of the base monolayer, h_m_; the peak maximum of fluorescence intensity, I(h_p_); and tail-to-monolayer intensity ratio, Φ_tm_. For relative comparison to monocultures, we divided those values obtained from co-culture biofilms by the corresponding values from the CB-D monoculture biofilms (Fig 4). Accordingly, a relative trend > 1 indicated an increase of the structural parameter (h_m_, I, or Φ_tm_) in co-culture as compared to monoculture. Whereas a relative trend < 1 indicated a decrease of the structural parameters (h_m_, I, or Φ_tm_) in co-culture compared to monoculture.

**Figure 4.**
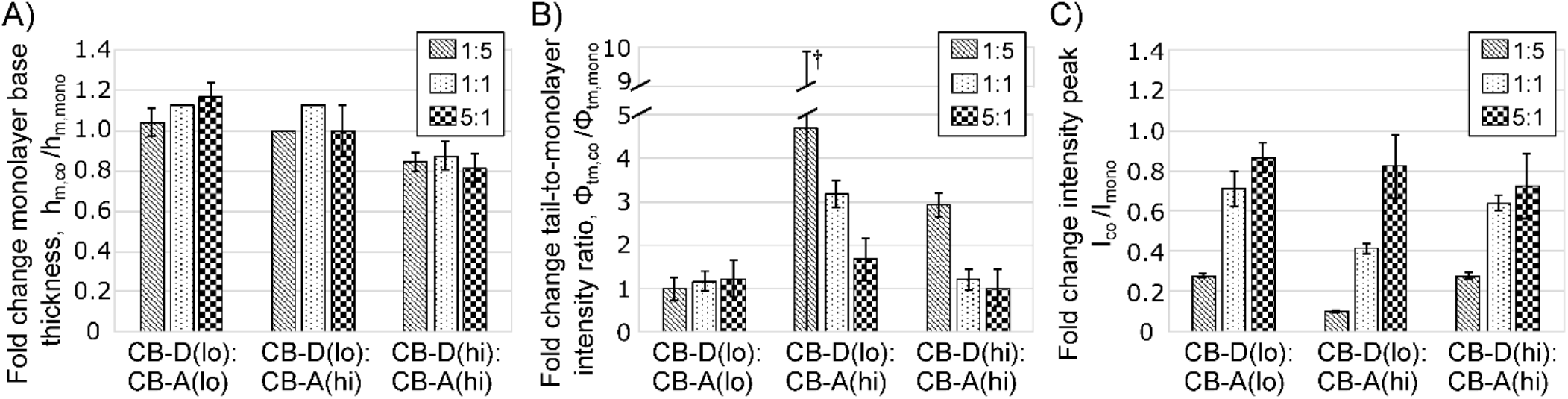
Structural characteristics of CB-D(lo) and CB-D(hi) in co-culture biofilms differ from those in monoculture biofilms. Fold-change was calculated for A) the mean monolayer base thickness, B) tail-to-monolayer intensity ratio, and C) fluorescence intensity peak **b**y extracting the fluorescence-labeled CB-D(lo) and CB-D(hi) cells from three replicates of co-culture biofilms with different inoculation ratios CB-D:CB-A for low, mixed, and high inducer pairs, respectively, and divided by the mean values of the corresponding CB-D(lo) and CB-D(hi) mean values from monoculture experiments. See Table S4 for plot data values and details on the calculation of uncertainty values. (^†^The high level of uncertainty is the result of one outlier sample.)

The peak width (h_m_) remained comparable between the co-cultures and the monocultures, regardless of seeding ratios and strain combination (Fig 4A). There was a slight increase in h_m_ for the low-inducer pair, and a slight decrease for the high-inducer pair. The monolayer thickness may be overestimated for high cell densities in the biofilm due to imperfect deconvolution of out-of-focus light in CSLM images, which can lead to the observed changes in the fold-change of the monolayer base. However, the absolute difference in thickness was ≤ 600 nm, which equates to a difference of one slice in the confocal z-stack, confirming that the cell monolayer with polar attachment was retained in all cases. In contrast, Φ_tm_ in co-cultures was equal to or larger than in monocultures (Fig 4B). This indicates that co-cultures exhibit larger relative representation of cells in protrusions (the tail of the fluorescence intensity curves) than in the monolayer base. Notably, while there is only minimal increase in Φ_tm_ of CB-D for low-inducer co-culture biofilms, the increase became more substantial with enhanced presence of CB-A(hi) during inoculation: For the high-inducer pair, we observed a substantial, 2.9-fold increase of Φ_tm_ of CB-D(hi) only for the 1:5 ratio, where the CB-A(hi) strain started as the abundant strain. For the mixed-inducer pairs, we observed substantial increase in Φ_tm_ of CB-D(lo), and this increase becomes larger with increasing initial abundance of the CB-A(hi) cells during inoculation. For the 1:5 seeding ratio where the CB-A(hi) strain begins as abundant, Φ_tm_ of the CB-D(lo) strain increased by 4.7-fold in co-cultures.

An increase in Φ_tm_ can be attained through either a decrease in biomass in the monolayer base, or an increase of biomass in the protrusions. We were able to rule out the first option based on the maximum peak intensity, I(h_p_) (Fig 4C). In all co-cultured pairs, the measured relative CB-D peak intensity is similar to or larger than the estimated value (estimated values for I_co_ / I_mono_ at 1:1 inoculation ratio based on monoculture growth volumes (Fig 1F) are: low inducer 0.48, mixed inducer 0.41, high inducer 0.54), suggesting that CB-D strains in co-culture do not lead to lower cell densities in the monolayer base due to intra-strain interaction. Hence, we conclude that the observed increase in Φ_tm_ is caused by increased biomass of CB-D cells in the protrusions. While the CB-D(lo) strain is unable to produce extensive protrusions on its own, when co-cultured with the high producer reciprocal strain CB-A(hi) a substantial increase in protrusion biomass occurs, irrespective of initial seeding density. The co-culture synergistic effect is also evident for the CB-D(hi) strain. Consequently, we postulate that in co-cultures, cells of divergent genotype may take advantage of and grow together with abundant, protrusion-forming cells, ultimately enhancing biomass in biofilm protrusions.

Our co-culture analysis is limited to the spatial architecture and height-dependent biomass of CB-D strains; an analogous analysis for the CB-A strains in co-culture is not possible due to small, but significant, inconsistencies in the imaging of the monolayer of mCherry fluorescent cells. These inconsistencies were on the order of the difference in values measured in Fig 4. Nonetheless, we were able to conduct quantitative analysis of the entire biofilm to investigate how pairwise co-culturing impacted overall biofilm biomass and that of each strain. We first measured the fraction of CB-D and CB-A strains in the co-cultured biofilms inoculated at a 1:1 ratio (Fig 5A) and compared their estimated representation derived from biofilm volumes in monoculture experiments. We find that the final relative biomass of each strain varies across the different co-cultured pairs in a manner consistent with expected values from monocultures. This finding indicates that the representation of each strain in a mature (24 hr) biofilm can be explained by the difference in strain-specific biofilm production rates alone. Thus, the phage-mediated intrastrain interaction does not significantly alter the relative representation of each strain within measurable limits. In contrast, the absolute biofilm biomass is significantly impacted by the monoculture interaction. The biomass of co-cultured biofilms is larger than the volume of monoculture biofilms of either strain (Fig 5B). Thus, phage-mediated interactions impact biofilm development by increasing biovolume, but do not influence the relative abundance of either strain.

**Figure 5.**
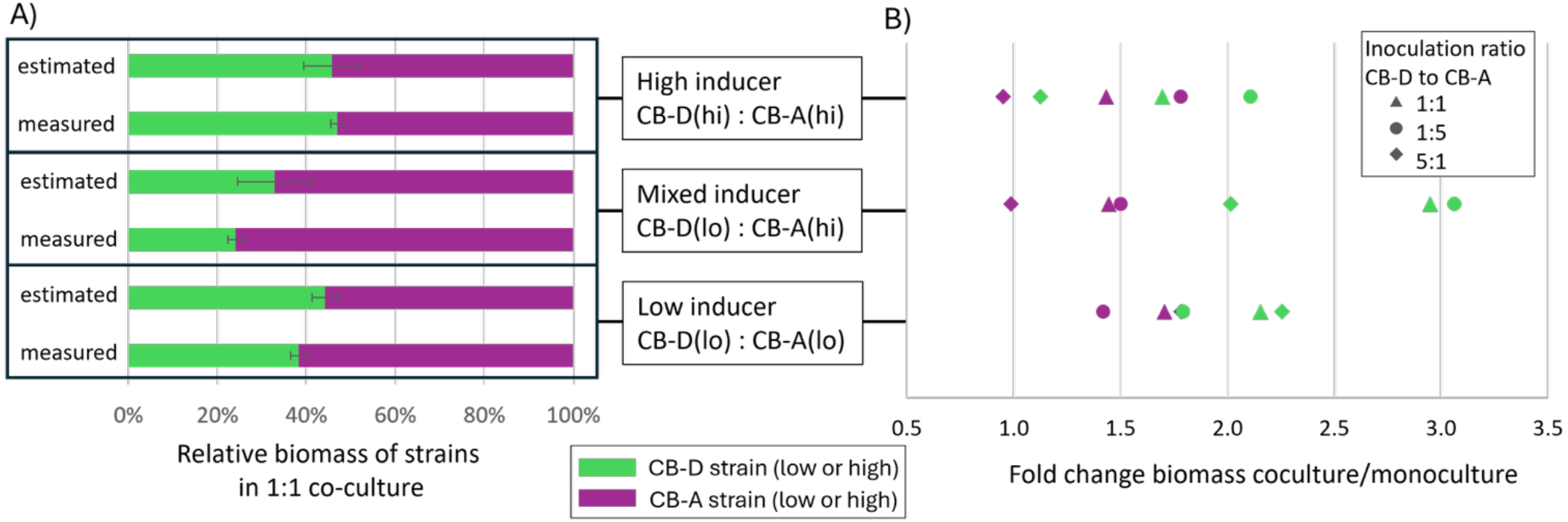
An increase in the absolute abundance of strain biomass, but no impact on relative abundance is evident in co-cultured biofilms. **(A)** The measured fractions of strains in co-cultured biofilms (high, mixed, and low inducer pairs) at 1:1 inoculation do not significantly differ from the estimated fractions derived from the ratio of monoculture biofilm production rates. **(B)** For all three co-cultured pairs and three inoculation ratios (1:1. 1:5, and 5:1), biomass fold change is obtained by the total biomass of co-culture biofilms divided by the biomass of the monoculture biofilm of either composing strain (green for CB-D and magenta for CB-A). In all cases, the biomass of co-culture biofilms is equal to or larger than either monoculture biofilm of the composing strains. Data for total biovolume and biovolume of CB-D and CB-A in co-culture biofilms are listed in supplemental Table S5.

### Co-culture biofilms show evidence of enhanced matrix production and intrastrain competition

Next, we assessed if production of biofilm matrix components was altered by induction state and co-culturing. To accomplish this, monoculture and co-culture biofilms were stained with fluorescent markers to observe major components of the biofilm matrix: extracellular DNA, matrix carbohydrates, and matrix proteins. For monocultures, we find that eDNA is dispersed through the biomatrix for all strains, forming a thin layer of eDNA at the biofilm base (Fig 6A). In addition, high inducer strains show localized high eDNA intensity within cells at or near the cell monolayer, likely indicative of dead cells with compromised cell walls. Increased levels of eDNA in high inducer strains is supported by an overall elevated level of total fluorescence (Fig 6C). We find similar signatures for CB-D(lo) albeit with lower overall intensity than high inducer strains. For co-cultured biofilms, lower eDNA fluorescence intensity tended to occur in low inducer co-cultures relative to either high inducer or mixed inducer co-culture pairs (Fig 6B, C). Enhanced eDNA intensity is visible throughout the monolayer and the protrusions for mixed and high inducer co-cultures; mixed inducer co-cultures show the greatest evidence of dead cells. Mixed and high inducer co-cultures show some cases of larger cell death than in monocultures, but not consistently (results for ratio 5:1 deviate) and in many cases the difference is not significant (Fig 6C).

**Figure 6.**
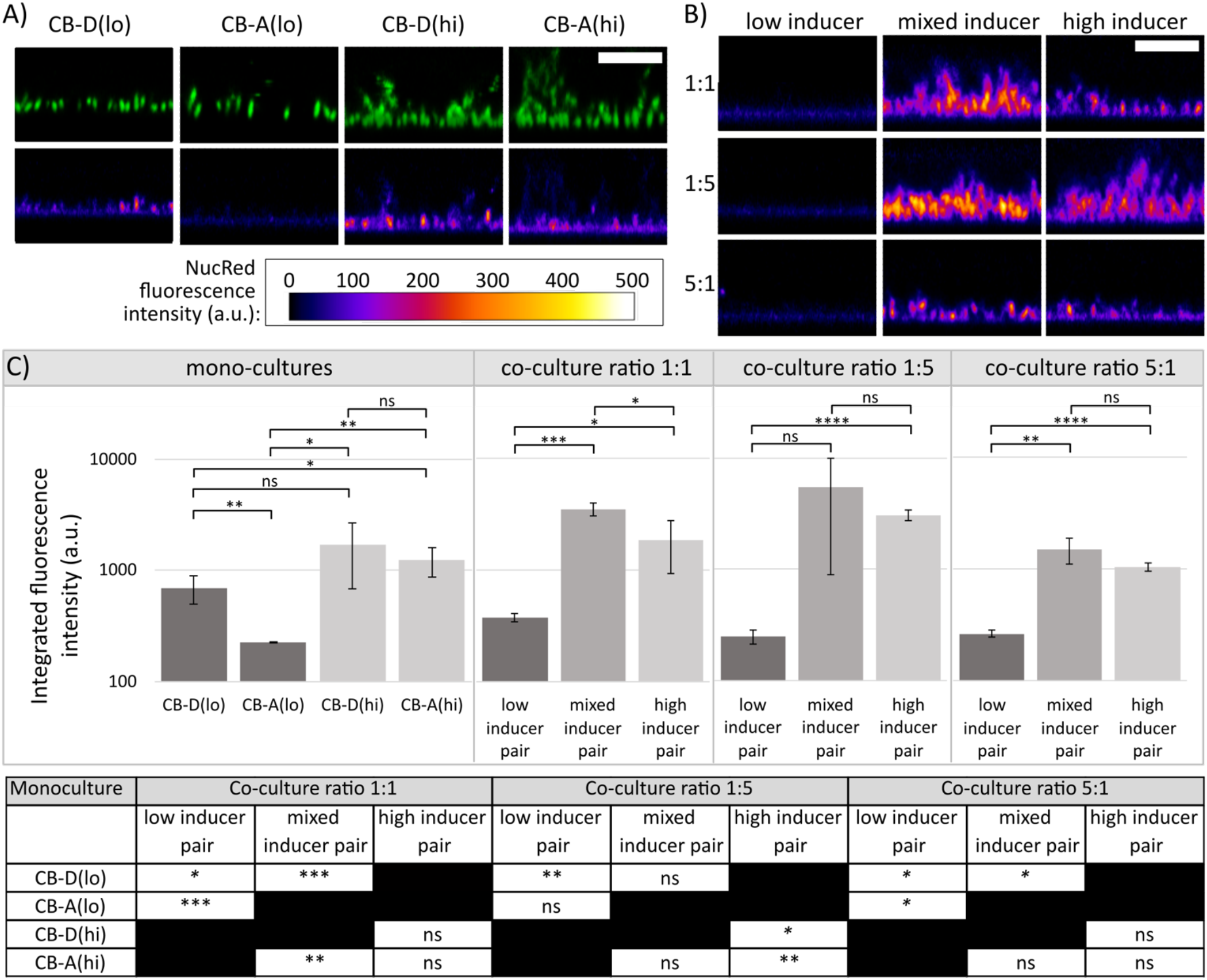
Evidence of dead cells and extracellular DNA within monoculture and co-culture biofilms. **A)** Side view of *S. pontiacus* monoculture biofilms grown for 24 h, displaying GFP signal from labeled cells (top row) and NucRed™ Dead 647 (bottom row), a cell-impermeable stain. **B)** Side view of *S. pontiacus* co-culture biofilms grown for 24 h stained with NucRed™ Dead 647 for three inoculation ratios and three pairs of co-cultured strains: low, mixed, and high inducer pairs (same as in Figures 3 and 4). Representative CSLM micrographs of labeled cells in co-culture biofilms from this experiment are shown in Figure 3. Scale bars (A and B): 10μm **C)** Integrated fluorescence intensity of NucRed™ Dead 647 of monoculture and co-culture biofilms in panels A and B. Bars represent mean values and standard deviations for three replicate Z-stack images. Student’s t-tests were performed to determine statistically significant differences between means of monoculture and co-culture biofilms, respectively (noted inside the plots), and across means of monocultures and corresponding co-culture pairings (noted in table below plots). ns = not significant, p > 0.05; * p ≤ 0.05; ** p ≤ 0.01; *** p ≤ 0.001; **** p ≤ 0.0001. Data for NucRed fluorescence intensities in panel C are listed in supplemental Table S6.

In monocultures, high-inducer strains show high levels of bound lectin-dye conjugate Concanavalin A (ConA) throughout the biofilm, while low-inducer biofilms have limited ConA binding, revealing that matrix carbohydrates increase with higher induction rates (Fig 7A). In addition, ConA signal was higher in the mixed inducer co-culture biofilms relative to monoculture biofilms of either induction state (Figs 7B, C), which is consistent with the enhanced biovolume observed in these co-culture biofilms (Fig 5B). Notably, in co-cultures, a large proportion of the bound ConA appeared to be associated with or in the vicinity of CB-A(hi) cells, whereas little binding of ConA is detected near CB-D(lo) cells (Fig 7B), consistent with the observations for ConA in monoculture biofilms. Results of nonspecific protein staining parallel those seen with ConA: the amount of matrix protein was highest in the mixed inducer co-culture biofilms relative to the monoculture biofilms of either induction state (Fig S4).

**Figure 7.**
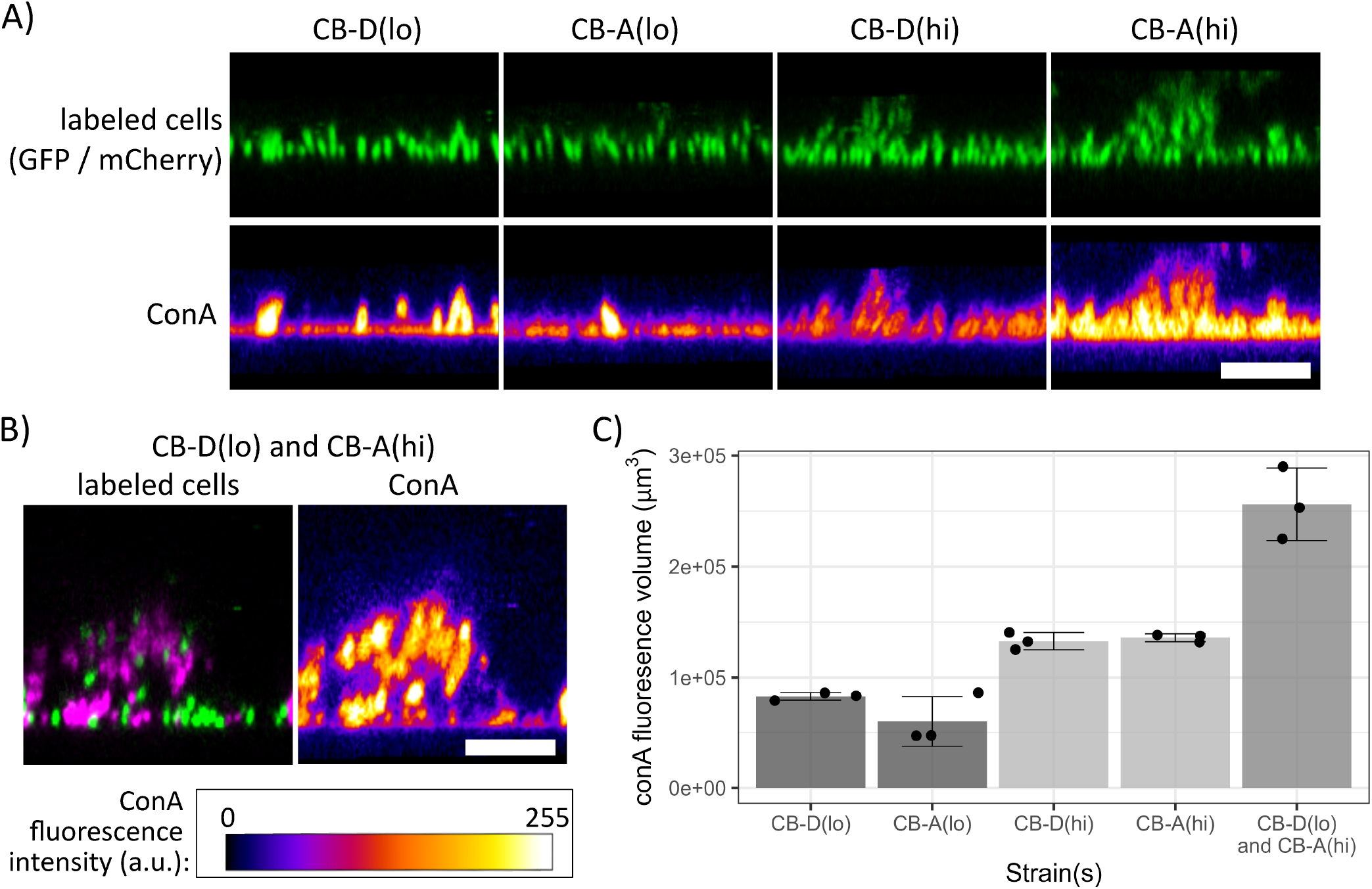
Staining with concanavalin A (ConA) shows an increase in biofilm matrix carbohydrates within high induction and co-culture biofilms. **A)** Side views of monoculture biofilms grown for 24 h showing fluorescence-labeled cells (top row) and ConA-CF405M dye conjugate staining (bottom row). *S. pontiacus* strains are constitutively labeled with GFP (CB-D(lo) and CB-D(hi)) or mCherry (CB-A(lo) and CB-A(hi)), both shown in green for better comparison. **B)** Side views of mixed-induction co-culture biofilm grown for 24 h (left) and stained with ConA-CF405M dye conjugate (right). *S. pontiacus* strains are constitutively labeled with GFP (CB-D(lo); green) and mCherry (CB-A(hi), magenta). Scale bars (A and B): 10 μm. **C)** Volume of ConA fluorescence within *S. pontiacus* biofilms. Bars represent mean values and standard deviations for three replicate Z-stack images (dots). Data for ConA fluorescence volumes in panel C are listed in supplemental Table S7.

### Co-culture biofilms contain higher levels of reactive oxygen species than monocultures

For many prophages, activation of the host SOS response in response to stress can lead to prophage induction (Butala et al., 2008; Nanda et al., 2015). Reactive oxygen species (ROS) such as superoxide (O_2_^-^), hydrogen peroxide (H_2_O_2_), and hydroxyl radicals (•OH) can induce an SOS response at high enough concentrations (Baharoglu and Mazel, 2014), which can lead to induction (Nanda et al., 2015). Oxidative stress is hypothesized to contribute to induction in our system, based on observations that D and A are mitomycin C-inducible and SOS response genes are upregulated with lytic infection (Basso et al., 2020). Therefore, we measured ROS levels within our biofilms using CellROX, an oxidative stress reporter that increases fluorescence intensity in the presence of ROS (Life Technologies, 2012). Studies have shown that this reagent can be used to quantify relative abundance of ROS within cells and is sensitive to the presence of O_2_^-^and •OH (Choi et al., 2015; McBee et al., 2017). ROS was detected within all biofilms, with stark differences between mono- and co-cultures (Fig 8). Renders of biofilms segmented into objects via cube segmentation reveal the highest levels of CellROX fluorescence were located near, but not on, the glass substrate within the surface-associated monolayer (Fig 8A). Distributions of the CellROX fluorescence intensity for each segmented object indicated statistically significant and strikingly higher levels of ROS within co-cultures compared to monoculture biofilms (Fig 8B). Additionally, within each biofilm, CellROX fluorescence of segmented objects was shown to positively correlate with the local density of cells, as measured by volume of mCherry fluorescence in the area around each object (Fig 8C). These data suggest that co-cultures experience higher levels of oxidative stress and that ROS may be playing an important role in competition within these biofilms.

**Figure 8.**
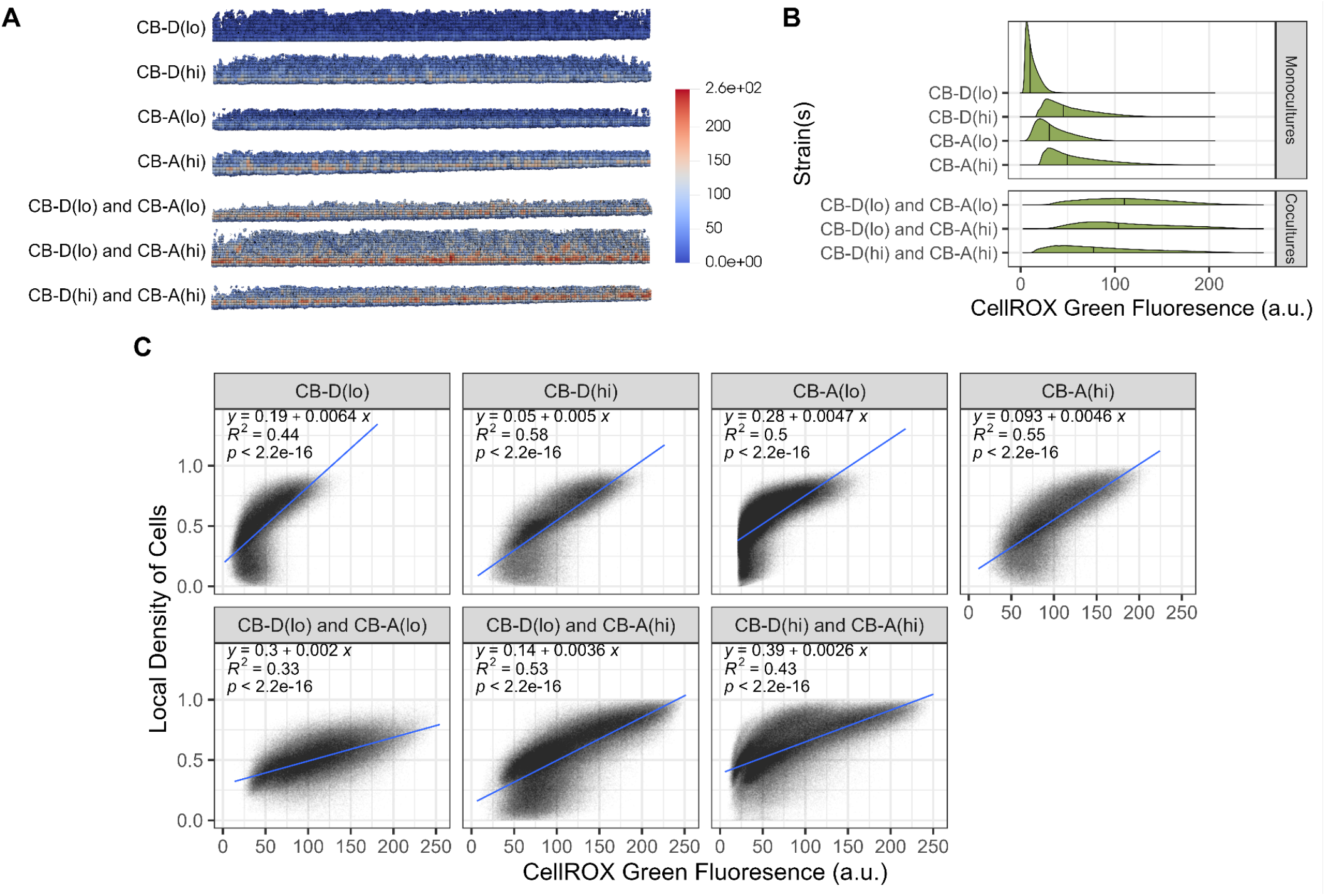
CellROX™ Green fluorescence within biofilms indicates higher levels of reactive oxygen species in multi-strain biofilms. **(A)** Side view of segmented biofilms rendered by ParaView. Renders depict the mean intensity of CellROX fluorescence and localization of segmented objects. **(B)** Frequency distribution of the average CellROX fluorescence (arbitrary units, a.u.) in biofilms by segmented object (n ≥ 42060 per Z-stack). Distributions for each biofilm include segmented objects from three Z-stack images per biofilm. Black lines depict median distribution values. Pairwise Wilcoxon tests indicated statistically significant differences between each distribution (p < 0.0001). **(C)** Correlations between local density of cells and average CellROX fluorescence by object in segmented biofilms. Local density is calculated as the occupied volume fraction of mCherry (monocultures) or SYTO41 (co-cultures) fluorescence in a sphere (radius: 12 voxels, 2.17 µm) around each segmented object. Linear regression lines (blue) were calculated using R.

## DISCUSSION

Phage-mediated intrastrain interactions are expected to be pervasive in dense microbial communities, where phages are abundant and often display narrow host specificity. While the ecological consequences of phage activity in spatially explicit systems remains poorly understood, there is increasing evidence that prophages play an important role in biofilm development and organization (e.g., Pires et al., 2021). By comparing monoculture and co-culture biofilms of *Sulfitobacter pontiacus* strains carrying either prophage type (A or D), we provide single cell-resolved information detailing how prophage induction state shapes biofilm architecture and matrix composition. We also show that reciprocal phage-host interactions can synergistically enhance biofilm growth rather than disrupt coexistence of competitive strains.

Our monoculture results corroborate previous work showing that prophage induction enhances biofilm formation (Carrolo et al., 2010; Nanda et al., 2015), while extending these findings by demonstrating structural differences in biofilm architecture that is correlated with induction state. High-induction strains consistently formed taller, rougher biofilms with pronounced three-dimensional protrusions, whereas low-induction strains produced flatter, more homogeneous biofilm topology. These architectural outcomes appear linked to the polar attachment of cells in *S. pontiacus* biofilms, with protrusions emerging from a basal monolayer in which cells maintain a vertical orientation. This contrasts with the smoother, dome-like biofilms of *Pseudomonas aeruginosa* and *Escherichia coli* (Klausen et al., 2003; Besharova et al., 2016) and the dome-shaped microcolonies formed by verticalization in *Vibrio cholerae* (Beroz et al., 2018; Hartmann et al., 2019, Drescher et al., 2016). The *S. pontiacus* biofilm initiation process resembles that of the alphaproteobacterium *Caulobacter crescentus*, although mature mushroom-shaped structures in *C. crescentus* arise from planarly organized cells (Entcheva-Dimitrov and Spormann, 2004; Berne et al., 2023), suggesting two biofilm forms distinguished by cell orientation relative to the surface: polar monolayers and planar mushroom structures. In *S. pontiacus*, phage-mediated lysis stabilizes pole-to-pole protrusions arising from the monolayer, thereby increasing total biofilm biomass. Taken together, these observations raise the intriguing possibility that prophage induction may actively influence the orientation of cells within protrusions, shaping the overall architecture of biofilms—a hypothesis that warrants further investigation.

Phage activity also proved central to co-culture outcomes. Reciprocal lysogens, which can kill one another through prophage induction (Basso et al., 2020), coexisted across all inoculation ratios and co-cultured pairs of strains. This result is striking given the expectation that strong local antagonism in dense systems often drives spatial assortment or competitive exclusion (Momeni et al., 2013; Nadell et al., 2016). As phage diffuse through biofilms (Winans et al., 2024), spatial assortment may be disrupted by the diffusion length of phage within these thin biofilm structures. Instead, we observed synergistic enhancement of biofilm growth, characterized by more frequent and larger protrusions. This effect was particularly pronounced when high- and low-induction strains were mixed, with low-induction cells visibly incorporated into protrusions dominated by high-induction counterparts.

Our findings on the spatial structure and biovolume reveal that phage-induced lysis can enhance biofilm formation in mixed inducer co-culture conditions. Co-cultures exhibited higher levels of extracellular DNA, carbohydrates, and proteins; all canonical contributors to biofilm stability and architecture (Carrolo et al., 2010; Nanda et al., 2015). Elevated reactive oxygen species (ROS) levels in co-cultures further support a role for phage-driven cell stress. While the source of ROS remains unclear, similar increases in oxidative stress have been linked to competitive interactions mediated by type VI secretion systems, antibiotic production, and phage attacks in other bacteria (Dong et al., 2015). ROS exposure can activate the bacterial SOS response, which in turn promotes prophage induction (Baharoglu and Mazel, 2014; Basso et al., 2020). Thus, ROS and prophage activity may form a positive feedback loop, amplifying induction in co-culture biofilms. Additional triggers such as plasmid conjugation (Baharoglu et al., 2010) or temperate phage infections themselves (Campoy et al., 2006) could further strengthen this feedback.

Although our confocal imaging approach has limitations, including the persistence of fluorescent signals in dead cells (Snapp, 2009) and the potential for genotype switching (Table S8), the overall trends we observed are robust. Importantly, prophage induction state correlates with plasmid content in *S. pontiacus* (Tuttle et al., 2022), indicating that biofilm development in this system is shaped by interactions among multiple mobile genetic elements.

Overall, the combined study of mono- and co-cultured biofilms reveals that phages, often thought of primarily as agents of mortality, can stabilize and enhance biofilm growth through mechanisms that combine antagonism with cooperation. This dual role embodied by prophages mirrors findings from other systems where temperate phages mediate both conflict and facilitation within microbial communities (e.g., Davies et al., 2016; Hargreaves et al., 2014). In our focal, ecologically relevant *S. pontiacus* system, mutually killing strains form stronger biofilms when coexisting in co-cultured conditions, demonstrating that population-level outcomes can emerge from the interplay of cell lysis, stress responses, and the corresponding matrix production. More broadly, our study adds to a growing recognition that phages shape microbial spatial organization not only by killing hosts but also by contributing to the stability and productivity of biofilm communities.

## METHODS

### Bacterial strains and culture conditions

Strains used in this study are listed in Table S9. *Sulfitobacter pontiacus* strains were routinely grown at 25°C in standard marine medium (SMM) amended with yeast extract and tryptone as previously described (Basso et al., 2020). Growth curves (Fig S5) were conducted using 10 ml SMM in 18 mm glass culture tubes and measured at OD_700_ to prevent bias in absorbance due to fluorescent protein expression (Hecht et al., 2016). *E. coli* strains harboring plasmids were routinely grown at 37°C in LB amended with 50 µg/ml kanamycin and 0.3 mM diaminopimelic acid (DAP). To investigate the cellular composition and structure of biofilms, we constructed a series of fluorescent strains of *S. pontiacus* CB-D(lo), CB-A(lo), CB-D(hi), and CB-A(hi) (Table S9). Insertion was performed using a modified Tn5 transposon for chromosomal insertion and increased stability in the absence of selective pressure. Plasmids and primers used in this study for the generation of strains are listed in Tables S10 and S11, respectively. Overlap extension PCR was performed to insert a *Bam*HI restriction site into the backbone of Tn5 transposon-containing plasmid pRL27 with KOD Xtreme Hot Start DNA Polymerase (Novagen Inc., USA) and the primers pRL27_OE_BamHI_Fw and pRL27_OE_BamHI_Rv. Thermocycling conditions were as follows: 2 min at 94°C, then 35 cycles of 10 s at 98°C, 30 s at 59°C, and 4.5 min at 68°C. Product was purified using a QIAquick PCR Purification Kit (QIAGEN, Germany), digested with *Bam*HI (Thermo Scientific, USA), purified again with the QIAquick PCR Purification Kit, and then ligated using T4 DNA Ligase (Promega, USA). Fluorescent versions of this plasmid were generated by first amplifying eGFP and mCherry genes (under control of the constitutive Alphaproteobacteria promoter A1/04/03) from the plasmids pBT211 and pBT277 (Zhao et al., 2013), respectively, using Platinum Taq (Invitrogen, USA) and the primers oJDR96_BamHI and oJDR97_BamHI. Thermocycling conditions were as follows: 2 min at 94°C, then 35 cycles of 15 s at 94°C, 30 s at 59°C, and 1.5 min at 72°C. Products were cut with *Bam*HI (Thermo Scientific, USA), pRL27-BamHI dephosphorylated with FastAP Alkaline Phosphatase (Thermo Scientific, USA), and cut DNA ligated with T4 DNA Ligase (Promega, USA). Plasmids were transformed into the chemically competent mating strain *E. coli* WM3064 (a DAP auxotroph; Saltikov and Newman, 2003). Plasmids containing fluorophores for tagging were delivered to *S. pontiacus* strains via biparental mating with *E. coli* WM3064 on an SMM agar plate. Transconjugants were enriched for and isolated on SMM amended with 50 µg/ml kanamycin (Km) in the absence of DAP. Tn5 insertional sites were identified via Sanger sequencing of arbitrary PCR products as previously described (Cude et al., 2012; O’Toole and Kolter, 1998). Assays of growth dynamics, phage production in the absence of prophage induction agents, and biofilm formation as described herein were performed on all strains to determine if Tn5 insertions occurred at neutral sites within strains (Figs S5-S7). Minimal to no differences in growth dynamics and phage production were seen between the original parent strains and their derivative fluorescent clones (Figs S5 and S6). Minor differences in biofilm formation were measured by a standard crystal violet biofilm assay (Fig S7). Following isolation of fluorophore-tagged *S. pontiacus* strains, subsequent experiments were conducted in the absence of Km.

### Confocal scanning laser microscopy (CSLM)

Strains were grown as single or multi-strain biofilms in 3 ml of SMM within 32 mm poly-D-lysine coated FluoroDishes (World Precision Instruments, USA). Dishes were seeded with ~10^7^ CFU/mL and for co-cultures, inoculation was performed at 5:1, 1:1, or 1:5 cell ratios. All dishes were incubated without agitation at 25°C for 24h. A subset of biofilms were then stained with 100 µg/ml Concanavalin A CF® Dye Conjugate (ConA) CF405M dye conjugate (lectin that selectively binds to α-mannopyranosyl and α-glucopyranosyl sugar moieties of glycoproteins and represents matrix carbohydrates; Biotium, USA), 3 ml FilmTracer™ SYPRO™ Ruby Biofilm Matrix Stain (nonspecific protein stain that interacts with primary amines of amino acids; Invitrogen, USA), or 2 drops/ml NucRed™ Dead 647 Ready Probes™ Reagent (cell impermeable DNA stain that binds to eDNA in the matrix as well as dead cells with compromised cell walls; Invitrogen, USA), followed by incubation for 30 min at room temperature (~25°C). For experiments measuring oxidative stress, biofilms were stained with 5 µM CellROX Green Reagent (Invitrogen, USA) and 5 µM SYTO41 (Invitrogen, USA) for 30 min and then fixed with 2.5% glutaraldehyde for 15 min. Planktonic cells and excess stain were removed from stained and unstained samples by gentle inversion and biofilms were washed three times with 3 ml SMM. Images were generated using an inverted Leica SP8 White Light Laser Confocal System at the University of Tennessee’s Advanced Microscopy and Imaging Center and visualized using Leica Application Suite X (LAS X) software (Leica Microsystems, USA). Z-stack images were taken using a 63X objective lens in 1024×1024 format with a line average of 3 and a LAS X system-optimized step size. Three replicate stacks were taken per sample, up to 100 images per stack, depending on biofilm sample thickness. Sequential scanning was used for all samples containing multiple fluorophores and hybrid (HyD) detectors were used to capture fluorescence. Microscope settings were consistent between compared samples unless otherwise noted.

### CSLM image analysis

Quantitative analysis of CSLM images was performed using BiofilmQ (version 0.2.2; Hartmann et al., 2021) for Figs 1, 5, 7C, 8, S1, S4C, and Tables S1, S2, S5, S7 and FIJI (version 1.53c, Schindelin et al., 2012) for Figs 2-4, 6, S2, S3, S4A-B, and Tables S3, S4, S6.

For data analyzed with BiofilmQ, image convolution, suppression of floating cells, and top-hat filtering were used to denoise images. Segmentation was performed using the otsu thresholding method (3-class with class 2 assigned to foreground, sensitivity = 0.5) and dissection using the cube method (cube side length = 6 vox [1.08 µm]) prior to calculation of parameters and plotting. Rendered 3D images of segmented biofilms were exported and visualized with ParaView (version 5.9.1; Kitware Inc., USA). The R programming language (version 3.6.0; R Core Team, 2017) was used for plotting with the Tidyverse collection of packages (Wickham et al., 2019) and ggridges (version 0.5.3; Wilke, 2021). Statistical tests were performed using ggpubr (version 0.2.5; Kassambara, 2020). For data analyzed with FIJI, the following image correction procedure was applied. Z-stacks of CSLM images were deconvoluted with the DeconvolutionLab2 plugin (Sage et al., 2017) using the Richardson-Lucy deconvolution algorithm over 4 iterations. The applied point spread function was created with the Born-Wolf model using the FIJI PSF generator plugin with the following settings: Refractive Index = 1, Accuracy = Good, NA = 1.2, Point Spread Function size = 50-pixel x 50-pixel x 20-pixel, Display = 8 bit. Suppression of floating cells was achieved by applying a vertical median filter with a range of 1 pixel. The vertical resolution of z-stacks was doubled through linear interpolation such that the distance between adjacent slices is half that of the original z-stack. To correct the tilting of samples during imaging, the biofilm stacks were brought into horizontal alignment using the TransformJ plugin (Meijering et al., 2001). For image analysis, the average fluorescence intensity, I(h), along the z-axis for each stack was obtained using the Plot Z-axis Profile plugin in FIJI. The z-stack slice with the maximum fluorescence intensity was designated as h_p_. The bottom of the biofilm, h_0_ (the glass substrate surface), was set by the lowest z-stack slice where the fluorescence intensity I(h) was at least 10% of the peak intensity of the stack: h_0_ was defined such that I(h_0_) = 0.1 I(h_p_). The thickness of the monolayer base was defined as h_m_=2(h_p_ - h_0_). The tail-to-monolayer intensity, Φ_tm_, which is a measure of the ratio of biomass between the protrusions and the monolayer of the biofilms (dark gray area divided by light grey area in Figure M1) was defined as 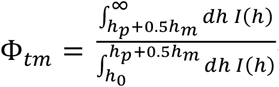.

**Figure M1.**
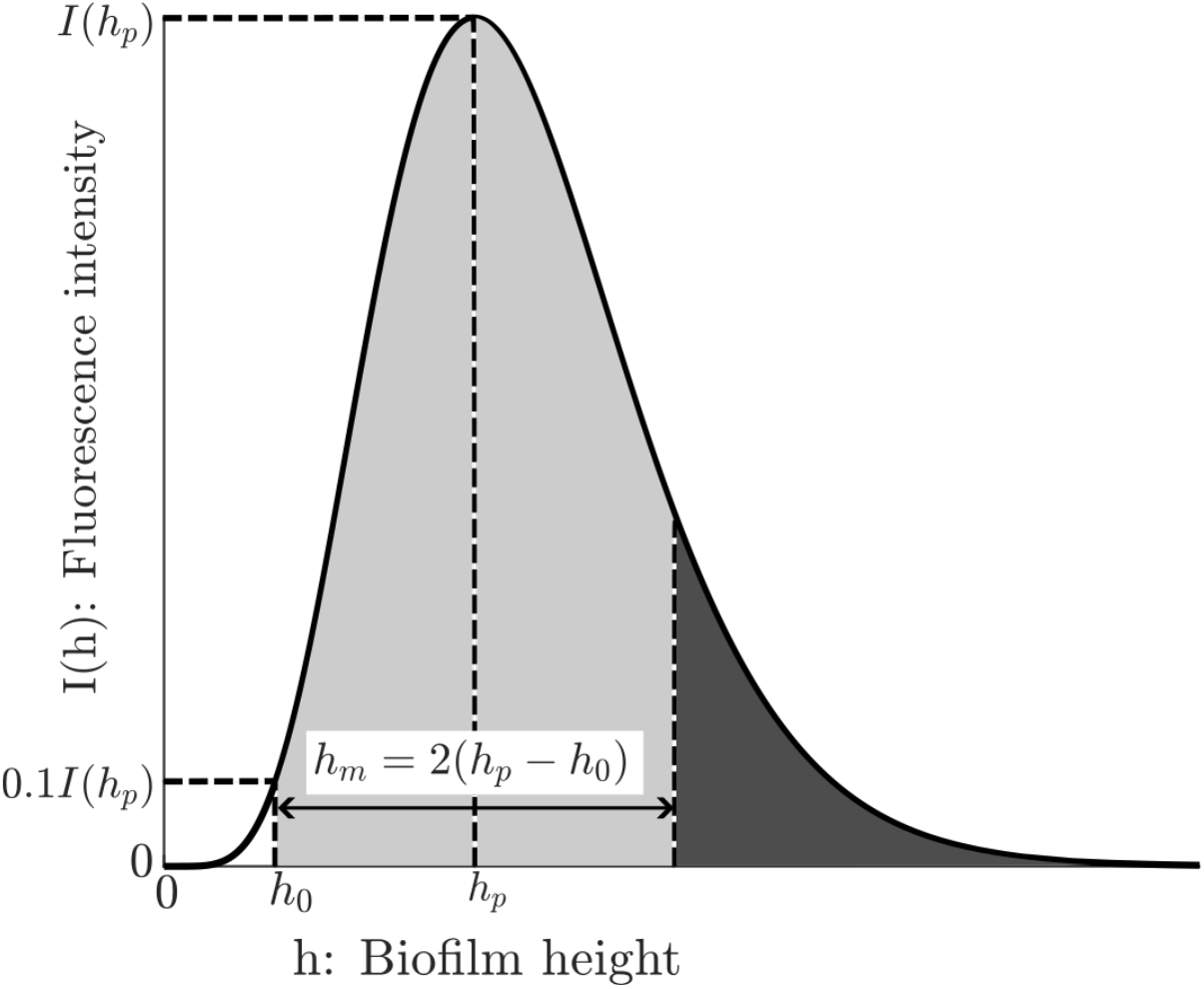
Schematic representation of height-dependent fluorescence intensity of imaged biofilms and relevant markers to describe biofilm structure and biomass distribution.

Fig M1 provides a schematic explaining the quantities. The shape and volume of protrusions in monoculture biofilms were obtained in FIJI by segmentation of the image-corrected sub-stack with biofilm heights at or above h_m_.

## ACKNOWLEDGEMENTS

The authors would like to thank Jennifer Morrell-Falvey for providing pBT211 and pBT277 courtesy of Boo Shan Tseng, Erik Zinser for providing *E. coli* WM3064, Barbara Klein for assistance in determining rates of lysogenic conversion among co-cultured strains, Jaydeep Kolape for CSLM training, Cameron Jackson for CSLM assistance, and Scott Retterer for discussions pertaining to fluorescent tagging of cells.

