## Supplemental Tables and Figures for "Prophage induction states drive structural and synergistic outcomes in marine bacterial biofilms"

**Table S1. Global properties of monoculture biofilms calculated by BiofilmQ.**

| Strain | Fluorophore | Rep | Biofilm_ Height (µm) | Biofilm MeanThickness (µm) | Biofilm Roughness (unitless) | Biofilm Volume ($\mu m^3$) |
|---|---|---|---|---|---|---|
| CB-D(lo) | GFP | 1 | 7.115 | 4.9176 | 0.15929 | 82939 |
| | | 2 | 7.1641 | 4.9121 | 0.15703 | 81159 |
| | | 3 | 7.3522 | 4.845 | 0.16002 | 82703 |
| | mCherry | 1 | 7.7777 | 4.8093 | 0.17148 | 72703 |
| | | 2 | 7.3129 | 4.9522 | 0.15816 | 83889 |
| | | 3 | 7.2325 | 5.1073 | 0.14903 | 90530 |
| CB-D(hi) | GFP | 1 | 14.4782 | 6.9824 | 0.25043 | 138372 |
| | | 2 | 16.0275 | 7.2617 | 0.26404 | 137835 |
| | | 3 | 16.931 | 7.445 | 0.29601 | 149541 |
| | mCherry | 1 | 19.5451 | 12.2934 | 0.3338 | 187615 |
| | | 2 | 18.5786 | 8.9471 | 0.34363 | 148547 |
| | | 3 | 19.7016 | 8.6659 | 0.34762 | 160103 |
| CB-A(hi) | GFP | 1 | 14.4227 | 5.5515 | 0.24915 | 111556 |
| | | 2 | 13.9555 | 5.936 | 0.24445 | 124198 |
| | | 3 | 15.041 | 6.677 | 0.22568 | 124406 |
| | mCherry | 1 | 13.8612 | 8.0194 | 0.26996 | 132316 |
| | | 2 | 19.8768 | 12.0093 | 0.36507 | 183946 |
| | | 3 | 19.6609 | 11.9869 | 0.36151 | 187164 |
| CB-A(lo) | GFP | 1 | 8.2498 | 5.594 | 0.16978 | 90426 |
| | | 2 | 8.2486 | 5.5289 | 0.1681 | 88890 |
| | | 3 | 8.1913 | 5.3083 | 0.17106 | 91235 |
| | mCherry | 1 | 7.4352 | 7.1592 | 0.12726 | 112835 |
| | | 2 | 9.0072 | 5.6436 | 0.15934 | 103434 |
| | | 3 | 7.1902 | 5.281 | 0.14831 | 94790 |

**Table S2. Means comparisons between GFP- and mCherry-fluorescent monoculture biofilms.**

| Property | Strain | Mean ± SD | | P-value[α] |
|---|---|---|---|---|
| | | GFP | mCherry | |
| Biofilm maximum height | CB-D(lo) | 7.21 ± 0.13 | 7.44 ± 0.29 | 0.30905 |
| | CB-D(hi) | 15.81 ± 1.24 | 19.28 ± 0.61 | 0.02402 |
| | CB-A(hi) | 14.47 ± 0.54 | 17.80 ± 3.41 | 0.23134 |
| | CB-A(lo) | 8.23 ± 0.03 | 7.88 ± 0.99 | 0.59909 |
| Biofilm average height | CB-D(lo) | 4.89 ± 0.04 | 4.96 ± 0.15 | 0.53494 |
| | CB-D(hi) | 7.23 ± 0.23 | 9.97 ± 2.02 | 0.14134 |
| | CB-A(hi) | 6.05 ± 0.57 | 10.67 ± 2.30 | 0.06587 |
| | CB-A(lo) | 5.48 ± 0.15 | 6.03 ± 1.00 | 0.43987 |
| Biofilm Roughness | CB-D(lo) | 0.159 ± 0.002 | 0.160 ± 0.011 | 0.91648 |
| | CB-D(hi) | 0.270 ± 0.023 | 0.342 ± 0.007 | 0.02572 |
| | CB-A(hi) | 0.240 ± 0.012 | 0.332 ± 0.054 | 0.09037 |
| | CB-A(lo) | 0.170 ± 0.001 | 0.145 ± 0.016 | 0.11866 |
| Biofilm Volume | CB-D(lo) | 8.23E+04 ± 9.67E+02 | 8.24E+04 ± 9.01E+03 | 0.98548 |
| | CB-D(hi) | 1.42E+05 ± 6.61E+03 | 1.65E+05 ± 2.01E+04 | 0.17088 |
| | CB-A(hi) | 1.20E+05 ± 7.36E+03 | 1.68E+05 ± 3.08E+04 | 0.10795 |
| | CB-A(lo) | 9.02E+04 ± 1.19E+03 | 1.04E+05 ± 9.03E+03 | 0.11982 |

[α] Means comparison performed in R using Student's t-tests

**Table S3. Biofilm structural characteristics of GFP-labeled monocultures after 24 h. Values reported here are (mean ± standard deviation) from 3 samples for each strain.**

| Strain | Thickness of monolayer base, $h_m$ (μm) | Tail-to-monolayer intensity ratio, $\Phi_{tm}$ | Peak fluorescence intensity $I(h_p)$ |
|---|---|---|---|
| CB-D(lo) | 2.39 ± 0.00 | 0.17 ± 0.02 | 116 ± 4 |
| CB-D(hi) | 3.18 ± 0.17 | 0.31 ± 0.03 | 212 ± 12 |
| CB-A(hi) | 2.79 ± 0.17 | 0.29 ± 0.05 | 190 ± 41 |
| CB-A(lo) | 2.79 ± 0.07 | 0.18 ± 0.04 | 119 ± 5 |

**Table S4: Structural characteristics of different CB-D inducer strains in co-cultures versus monoculture.**

| Co-cultured strains | Inoculation ratio | Relative monolayer base thickness[a] $h_{m,co} / h_{m,mono}$ | Relative tail-to-monolayer intensity ratio[a] $\Phi_{tm,co} / \Phi_{tm,mono}$ | Relative intensity peak[a] $I_{co} / I_{mono}$ |
|---|---|---|---|---|
| CB-D(hi) : CB-A(hi) (high-inducer) | 5:1 | 0.813 ± 0.07 | 1.012 ± 0.113 | 0.722± 0.161 |
| | 1:1 | 0.875 ± 0.072 | 1.208 ± 0.156 | 0.639± 0.035 |
| | 1:5 | 0.844 ± 0.046 | 2.925 ± 0.357 | 0.276± 0.016 |
| CB-D(lo) : CB-A(hi) (mixed-inducer) | 5:1 | 1.000 ± 0.125 | 1.674 ± 0.485 | 0.821± 0.159 |
| | 1:1 | 1.125 ± 0.0 | 3.180 ± 0.301 | 0.412± 0.024 |
| | 1:5 | 1.000 ± 0.0 | 4.700 ± 5.181* | 0.101± 0.006 |
| CB-D(lo) : CB-A(lo) (low-inducer) | 5:1 | 1.167 ± 0.072 | 1.230 ± 0.428 | 0.862 ± 0.075 |
| | 1:1 | 1.125 ± 0.00 | 1.171 ± 0.229 | 0.709 ± 0.086 |
| | 1:5 | 1.042 ± 0.072 | 0.993 ± 0.269 | 0.277 ± 0.013 |

[a] Mean monolayer base thickness, tail-to-monolayer intensity ratio, and fluorescence intensity peak were extracted for the fluorescence-labeled CB-D cells in co-culture biofilms, and values were divided by the corresponding mean values from monoculture experiments with the CB-D strain. The uncertainties were calculated as the product of the mean and the square root of the sum of squares of the coefficients of variation. So, if z = x/y. Then, $\delta z = z\sqrt{(\frac{\delta x}{x})^2 + (\frac{\delta y}{y})^2}$, where $\delta x$ and $\delta y$ are the standard deviations in the measurements of x and y. *The high level of uncertainty is the result of one outlier sample image.

**Table S5: Biovolume of strains in co-culture biofilms calculated by BiofilmQ.**

| Ratio$^{\alpha}$ | Strains | Replicate | GFP volume (µm$^3$) | mCherry volume (µm$^3$) | Total cell volume$^{\beta}$ (µm$^3$) | GFP volume relative to total cell volume | mCherry volume relative to total cell volume |
|---|---|---|---|---|---|---|---|
| 5:1 | CB-D(lo) and CB-A(lo) | 1 | 84709 | 41759 | 126468 | 66.98% | 33.02% |
| 5:1 | CB-D(lo) and CB-A(lo) | 2 | 90033 | 116018 | 206051 | 43.69% | 56.31% |
| 5:1 | CB-D(lo) and CB-A(lo) | 3 | 92173 | 131959 | 224132 | 41.12% | 58.88% |
| 1:1 | CB-D(lo) and CB-A(lo) | 1 | 63483 | 98123 | 161606 | 39.28% | 60.72% |
| 1:1 | CB-D(lo) and CB-A(lo) | 2 | 67422 | 119031 | 186453 | 36.16% | 63.84% |
| 1:1 | CB-D(lo) and CB-A(lo) | 3 | 72578 | 111393 | 183971 | 39.45% | 60.55% |
| 1:5 | CB-D(lo) and CB-A(lo) | 1 | 21683 | 123556 | 145238 | 14.93% | 85.07% |
| 1:5 | CB-D(lo) and CB-A(lo) | 2 | 22208 | 127384 | 149591 | 14.85% | 85.15% |
| 1:5 | CB-D(lo) and CB-A(lo) | 3 | 22631 | 124875 | 147507 | 15.34% | 84.66% |
| 5:1 | CB-D(lo) and CB-A(hi) | 1 | 66387 | 73733 | 140120 | 47.38% | 52.62% |
| 5:1 | CB-D(lo) and CB-A(hi) | 2 | 78322 | 87388 | 165709 | 47.26% | 52.74% |
| 5:1 | CB-D(lo) and CB-A(hi) | 3 | 92605 | 99067 | 191672 | 48.31% | 51.69% |
| 1:1 | CB-D(lo) and CB-A(hi) | 1 | 52317 | 184721 | 237038 | 22.07% | 77.93% |
| 1:1 | CB-D(lo) and CB-A(hi) | 2 | 58418 | 173749 | 232167 | 25.16% | 74.84% |
| 1:1 | CB-D(lo) and CB-A(hi) | 3 | 64524 | 194446 | 258970 | 24.92% | 75.08% |
| 1:5 | CB-D(lo) and CB-A(hi) | 1 | 16016 | 181084 | 197100 | 8.13% | 91.87% |
| 1:5 | CB-D(lo) and CB-A(hi) | 2 | 32679 | 352351 | 385029 | 8.49% | 91.51% |

| | | | | | | | |
|---|---|---|---|---|---|---|---|
| 1:5 | CB-D(lo) and CB-A(hi) | 3 | 10512 | 163453 | 173965 | 6.04% | 93.96% |
| 5:1 | CB-D(hi) and CB-A(hi) | 1 | 95609 | 41247 | 136856 | 69.86% | 30.14% |
| 5:1 | CB-D(hi) and CB-A(hi) | 2 | 112350 | 47402 | 159752 | 70.33% | 29.67% |
| 5:1 | CB-D(hi) and CB-A(hi) | 3 | 122942 | 59746 | 182688 | 67.30% | 32.70% |
| 1:1 | CB-D(hi) and CB-A(hi) | 1 | 113161 | 133548 | 246708 | 45.87% | 54.13% |
| 1:1 | CB-D(hi) and CB-A(hi) | 2 | 110300 | 127355 | 237655 | 46.41% | 53.59% |
| 1:1 | CB-D(hi) and CB-A(hi) | 3 | 115640 | 122369 | 238009 | 48.59% | 51.41% |
| 1:5 | CB-D(hi) and CB-A(hi) | 1 | 88409 | 194132 | 282541 | 31.29% | 68.71% |
| 1:5 | CB-D(hi) and CB-A(hi) | 2 | 86567 | 216673 | 303240 | 28.55% | 71.45% |
| 1:5 | CB-D(hi) and CB-A(hi) | 3 | 89625 | 222570 | 312195 | 28.71% | 71.29% |

$^{\alpha}$ Ratio of GFP fluorescent CB-D(lo) or CB-D(hi) to mCherry-fluorescent CB-A(lo) or CB-A(hi) added at initial inoculation.

$^{\beta}$ Sum of GFP and mCherry fluorescence volumes.

**Table S6: NucRed fluorescence intensity for mono-culture and co-culture biofilms**

| Strain(s) | Inoculation ratio | Replicate | Integrated NucRed intensity (a.u.) | Mean intensity ± SD |
|---|---|---|---|---|
| CB-D(lo) | N/A | 1 | 888 | 688 ± 195 |
| | | 2 | 679 | |
| | | 3 | 497 | |
| CB-A(lo) | N/A | 1 | 220 | 224 ± 3 |
| | | 2 | 227 | |
| | | 3 | 224 | |
| CB-D(hi) | N/A | 1 | 2761 | 1687 ± 1005 |
| | | 2 | 1531 | |
| | | 3 | 768 | |
| CB-A(hi) | N/A | 1 | 1648 | 1239 ± 362 |
| | | 2 | 1106 | |
| | | 3 | 963 | |
| CB-D(lo) and CB-A(lo) | 1:1 | 1 | 357 | 375 ± 29 |
| | | 2 | 408 | |
| | | 3 | 360 | |
| CB-D(lo) and CB-A(lo) | 1:5 | 1 | 290 | 250 ± 35 |
| | | 2 | 229 | |
| | | 3 | 231 | |
| CB-D(lo) and CB-A(lo) | 5:1 | 1 | 286 | 266 ± 20 |
| | | 2 | 247 | |
| | | 3 | 264 | |
| CB-D(lo) and CB-A(hi) | 1:1 | 1 | 4111 | 3530 ± 508 |
| | | 2 | 3314 | |
| | | 3 | 3166 | |
| CB-D(lo) and CB-A(hi) | 1:5 | 1 | 5149 | 5456 ± 4551 |
| | | 2 | 10153 | |
| | | 3 | 1065 | |
| CB-D(lo) and CB-A(hi) | 5:1 | 1 | 1781 | 1498 ± 377 |
| | | 2 | 1642 | |
| | | 3 | 1070 | |
| CB-D(hi) and CB-A(hi) | 1:1 | 1 | 2848 | 1840 ± 909 |
| | | 2 | 1592 | |
| | | 3 | 1080 | |
| CB-D(hi) and CB-A(hi) | 1:5 | 1 | 3324 | 3095 ± 326 |
| | | 2 | 2722 | |
| | | 3 | 3238 | |
| | 5:1 | 1 | 963 | 1045 ± 85 |

| | | | |
|---|---|---|---|
| CB-D(hi) and CB-A(hi) | | 2 | 1040 |
| | | 3 | 1132 |

**Table S7: ConA fluorescence intensity for mono-culture and co-culture biofilms**

| Strain(s) | Replicate | conA fluorescence volume ($\mu m^3$) | Mean volume ± SD |
|---|---|---|---|
| CB-D(lo) | 1 | 7.94E+04 | 8.30E+04 ± 3.46E+03 |
| | 2 | 8.36E+04 | |
| | 3 | 8.62E+04 | |
| CB-A(lo) | 1 | 8.64E+04 | 6.02E+04 ± 2.27E+04 |
| | 2 | 4.72E+04 | |
| | 3 | 4.70E+04 | |
| CB-D(hi) | 1 | 1.25E+05 | 1.33E+05 ± 7.73E+03 |
| | 2 | 1.32E+05 | |
| | 3 | 1.41E+05 | |
| CB-A(hi) | 1 | 1.38E+05 | 1.36E+05 ± 3.55E+03 |
| | 2 | 1.32E+05 | |
| | 3 | 1.38E+05 | |
| CB-D(lo) and CB-A(hi) | 1 | 2.53E+05 | 2.56E+05 ± 3.26E+04 |
| | 2 | 2.90E+05 | |
| | 3 | 2.25E+05 | |

**Table S8. Prophage genotype screening of GFP-Km$^R$ colonies plated from liquid co-cultures of CB-D Δ(2,4) and CB-A Δ(2,4) GFP after 4 days growth.** As infections by the temperate phages φ-D and φ-A lead to switching between prophage genotypes in a subpopulation of lytically infected lysogens (Basso et al., 2020), we sought to determine the potential rate of this conversion. To ensure high rates of infection and establish a potential maximum conversion rate, we co-cultured two strains exhibiting high rates of spontaneous prophage induction, CB-D(hi) and CB-A(hi) GFP (Figure S3). These strains were inoculated at a 1:1 ratio and grown planktonically for 4 days, prior to plating on media amended with Km. PCR screening of Km-resistant colonies (all of which possessed φ-A at inoculation) revealed that 13.3% underwent prophage genotype conversion, now possessing only φ-D (Table S6). An additional 13.3% possessed both φ-A and φ-D DNA, a state previously observed to be transient and unstable over time (Basso et al., 2020). These data indicate that prophage genotype conversion occurs in a relatively small proportion of the infected population. Thus, it is important to note that within subsequent experiments in this study involving co-cultures, the respective fluorophores observed represent the progeny of strains used as inoculum and are not necessarily reflective of the prophage genotype within these cells.

| **PCR screen result** | **Colonies screened** | **Percent of screened** |
|---|---|---|
| φ-A+ only | 22 | 73.3% |
| φ-D+ only | 4 | 13.3% |
| φ-A+ and φ-D+ | 4 | 13.3% |
| **Total** | **30** | **100%** |

**Table S9. Strains used in this study**

| Species | Strain | Description | Tn5 insertion site (locus tag for accession GCA_000735125.2) | Source/reference |
|---|---|---|---|---|
| *Sulfitobacter pontiacus* | CB-D(lo) | Also known as strain CB2047, isolated from an *Emiliania huxleyi* phytoplankton bloom in Raunefjorden, Norway. Possesses four plasmids: pSpoCB-1, pSpoCB-2, pSpoCB-3, and pSpoCB-4. | | (Ankrah et al., 2014) |
| | CB-D (lo) GFP | Constitutively fluorescent strain derived from CB-D via pRL27-GFP insertion. | hypothetical protein (IV89_002340) | This study |
| | CB-D (lo) mCherry | Constitutively fluorescent strain derived from CB-D via pRL27-mCherry insertion. | intergenic region (between IV89_001325 and IV89_001326) | This study |
| | CB-A(lo) | Derivative of CB-D from superinfection with φ-A. Possesses four plasmids: pSpoCB-1, pSpoCB-2, pSpoCB-3, and pSpoCB-4. | | (Tuttle et al., 2022) |
| | CB-A (lo) GFP | Constitutively fluorescent strain derived from CB-A via pRL27-GFP insertion. | purine nucleoside phosphorylase DeoD-type (IV89_002872) | This study |
| | CB-A (lo) mCherry | Constitutively fluorescent strain derived from CB-A via pRL27-mCherry insertion. | esterase (IV89_002996) | This study |
| | CB-D (hi) | Derivative of CB-A from superinfection with φ-D. Possesses two plasmids: pSpoCB-1 and pSpoCB-3. | | (Tuttle et al., 2022) |
| | CB-D (hi) GFP | Constitutively fluorescent strain derived from CB-D Δ(2,4) via pRL27-GFP insertion. | hypothetical protein (IV89_003033) | This study |
| | CB-D (hi) mCherry | Constitutively fluorescent strain derived from CB-D Δ(2,4) via pRL27-mCherry insertion. | intergenic region (between IV89_000082 and IV89_000083) | This study |
| | CB-A (hi) | Generated from superinfection of CB-D with φ-A. Possesses two plasmids: pSpoCB-1 and pSpoCB-3. | | (Basso et al., 2020) |
| | CB-A (hi) GFP | Constitutively fluorescent strain derived from CB-A Δ(2,4) via pRL27-GFP insertion. | hypothetical protein (IV89_001984) | This study |
| | CB-A (hi) mCherry | Constitutively fluorescent strain derived from CB-A Δ(2,4) via pRL27-mCherry insertion. | TonB-dependent siderophore receptor (IV89_000025) | This study |
| *Escherichia coli* | WM3064 | Mating strain, DAP auxotroph. | | (Saltikov and Newman, 2003) |

**Table S10. Plasmids used in this study.**

| Plasmid | Description | Source/reference |
|---|---|---|
| pBT211 | MiniTn7 transposon containing eGFP expressed under the A1/04/03 promoter, $Gm^R$. | (Zhao et al., 2013) |
| pBT277 | MiniTn7 transposon containing mCherry expressed under the A1/04/03 promoter, $Gm^R$. | (Zhao et al., 2013) |
| pRL27 | Tn5 transposon delivery vector, $Km^R$, *ori*R6K. | (Larsen et al., 2002) |
| pRL27-*Bam*HI | Tn5 transposon delivery vector containing a *Bam*HI site, $Km^R$, *ori*R6K. | This study |
| pRL27-GFP | Tn5 transposon delivery vector containing eGFP expressed under the A1/04/03 promoter, $Km^R$, *ori*R6K. | This study |
| pRL27-mCherry | Tn5 transposon delivery vector containing mCherry expressed under the A1/04/03 promoter, $Km^R$, *ori*R6K. | This study |

Km = kanamycin, Gm = Gentamicin

**Table S11. Oligonucleotides used in this study.**

| Primer name | Sequence (5' to 3')[a] | Purpose/target | Reference/ source |
|---|---|---|---|
| pRL27_OE_BamHI_Fw | GAT CGA GGA TCC CCC GTC AAG TCA GCG TAA TG | OE-PCR of pRL27 to construct pRL27-*Bam*HI. | This study |
| pRL27_OE_BamHI_Rv | GTG ATA GGA TCC ACG GCG GCT TTG TTG AAT AA | OE-PCR of pRL27 to construct pRL27-*Bam*HI. | This study |
| oJDR96_BamHI | ACG ATC GGA TCC ATC CTG AAA ATT TAT CAA AAA GAG TGT TGA C | Amplification of fluorophores off pBT211 and pBT277 flanked by *Bam*HI sites. Modified from (Rich, 2016). | This study |
| oJDR97_BamHI | ATT CGC GGA TCC ATC CTT ACT TGT ACA GCT CGT CCA TGC | Amplification of fluorophores off pBT211 and pBT277 flanked by *Bam*HI sites. Modified from (Rich, 2016). | This study |
| ARB6 | GGC CAC GCG TCG ACT AGT ACN NNN NNN NNN ACG CC | Identification of Tn5 insertion sites via arbitrary PCR. | (Cude et al., 2012) |
| ARB2 | GGC CAC GCG TCG ACT AGT AC | Identification of Tn5 insertion sites via arbitrary PCR. | (O'Toole and Kolter, 1998) |
| TNPR17In | CGT TAC ATC CCT GGC TTG TT | Confirmation of Tn5 transposon presence. | (Cude et al., 2012) |
| TNPR17Out | AAC AAG CCA GGG ATG TAA CG | Identification of Tn5 insertion sites via arbitrary PCR. | (Cude et al., 2012) |
| TNPR17Nest | CTG ACA TGG GGG GGT ACC | Identification of Tn5 insertion sites via arbitrary PCR. | (Cude et al., 2012) |
| TNPR13In | TCG TGA AGA AGG TGT TGC TG | Confirmation of Tn5 transposon presence. | (Cude et al., 2012) |

| | | | |
|---|---|---|---|
| TNPR13Out | CAG CAA CAC CTT CTT CAC GA | Identification of Tn5 insertion sites via arbitrary PCR. | (Cude et al., 2012) |
| TNPR13Nest | CTA GAG TCG ACC TGC AGG CAT | Identification of Tn5 insertion sites via arbitrary PCR. | (Cude et al., 2012) |
| 2047PP1.for | TAT TCA TAG CGA GGC GCA GT | Detection of φ-D presence. | (Basso et al., 2020) |
| 2047PP1.rev | ATA CCT GCC CAA CGT CAC AG | Detection of φ-D presence. | (Basso et al., 2020) |
| 2047A-C.for | CCC ATG TGT ATG TCG CCT CT | Detection of φ-A presence. | (Basso et al., 2020) |
| 2047A-C.rev | CAG CGT TGA AAA AGG CTC TG | Detection of φ-A presence. | (Basso et al., 2020) |

[α] Underlined nucleotides indicate *Bam*HI recognition sites

### Supplemental Figures

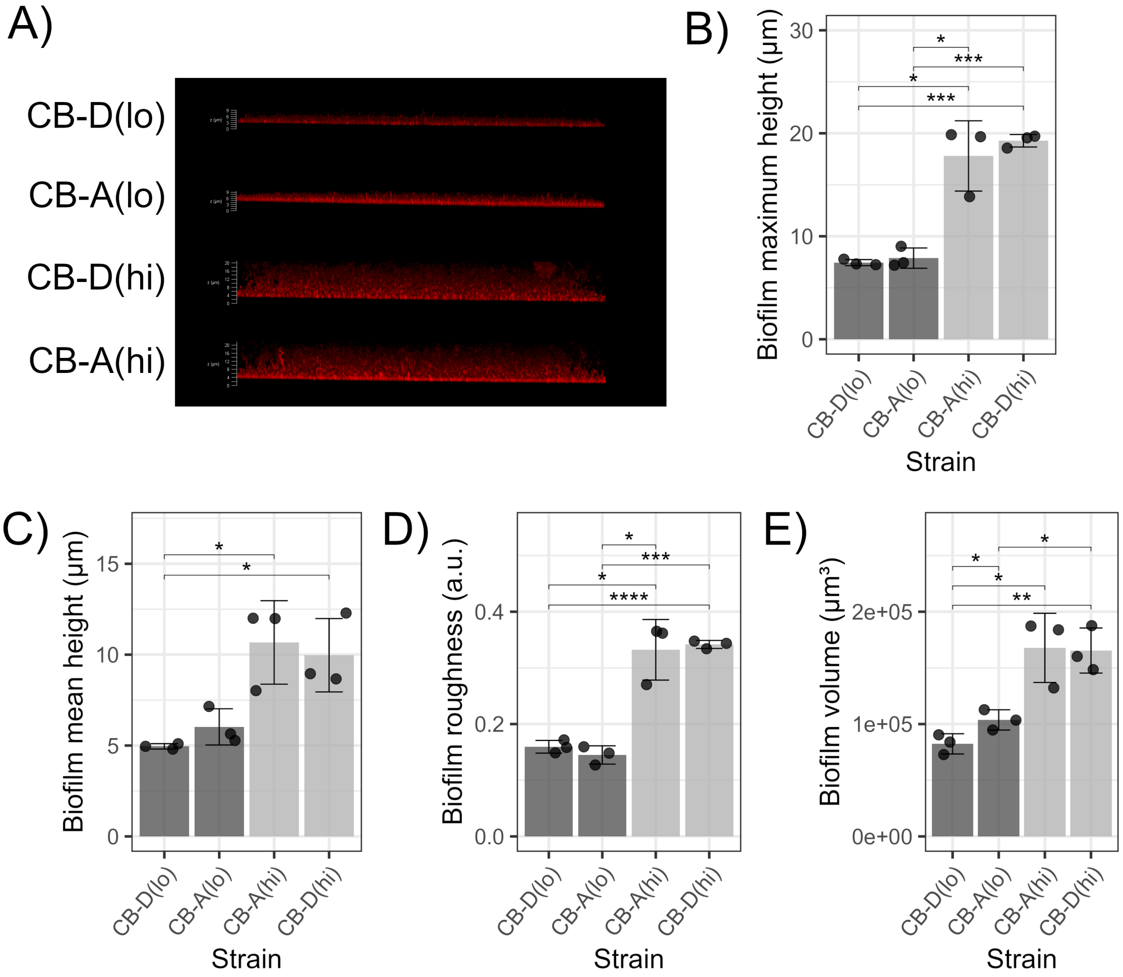

**Figure S1. Monoculture biofilms of mCherry-fluorescent strains exhibit similar characteristics to GFP-fluorescent biofilms. (A)** Side views of representative 3D CSLM micrographs of mCherry-fluorescent strains grown for 24 h as monoculture biofilms. **(B)** Maximum height of monoculture biofilms, defined as the height of z coordinates in the range [1%, 99%]. **(C)** Mean height of monoculture biofilms, defined as the mean height of each vertical stack of objects from cube segmentation of images. **(D)** Roughness coefficient of monoculture biofilms. **E)** Total volume of monoculture biofilms as measured by volume of mCherry fluorescence for images of lateral size 185 µm x 185 µm. For panels C-F points represent replicates from Z-stack images, columns represent the mean of all replicates (n = 3), and error bars depict standard deviation. Student's t-tests were performed to determine statistically significant differences between means (* $p \leq 0.05$; ** $p \leq 0.01$; *** $p \leq 0.001$; **** $p \leq 0.0001$). Data for biofilm properties measured in this experiment are available in Table S1 and S2.

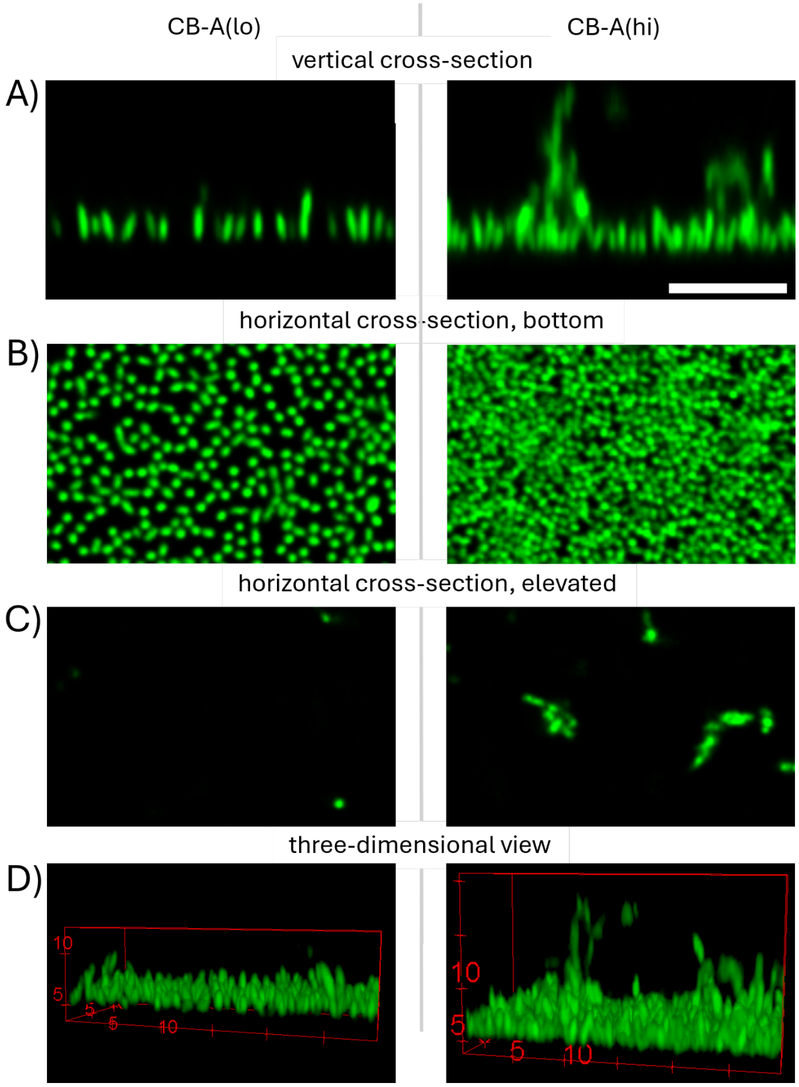

**Figure S2. Spatial structure of CB-A(lo) and CB-A(hi).** Representative confocal microscopy images of monoculture biofilms grown from CB-A(lo) and CB-A(hi) strains for 24 hours are shown as cross-sections from a A) vertical cut, B) horizontal cuts through the monoloayer (at $h_p$) and C) 6 μm above $h_p$, and D) a three-dimensional view.

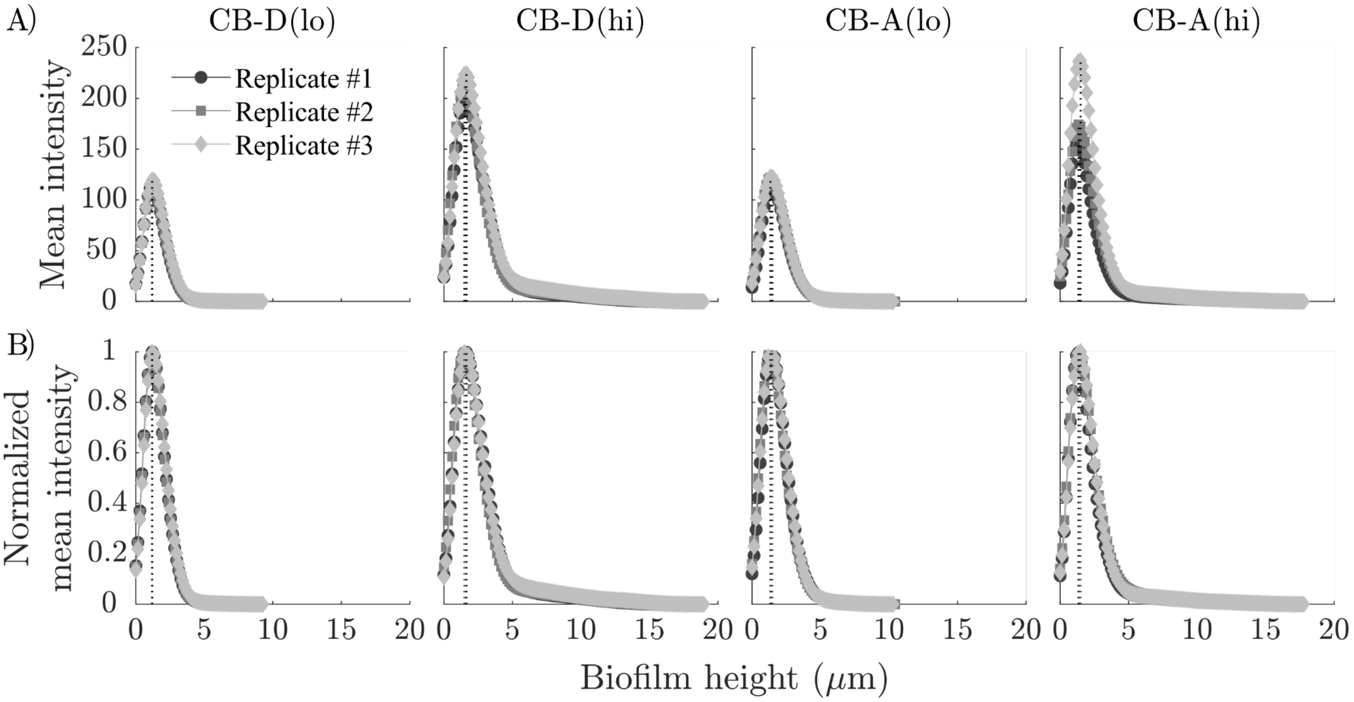

**Figure S3. Height-dependent fluorescence intensity of monoculture biofilms of GFP-fluorescent strains. (A)** Mean fluorescence intensity and **(B)** normalized mean fluorescence intensity as a function of biofilm height, where height h=0 corresponds to $h_0$ (see Fig M1 for reference). Each graph depicts curves of three representative samples. Vertical lines indicate the biofilm height at peak intensity, $h_p$, for each sample.

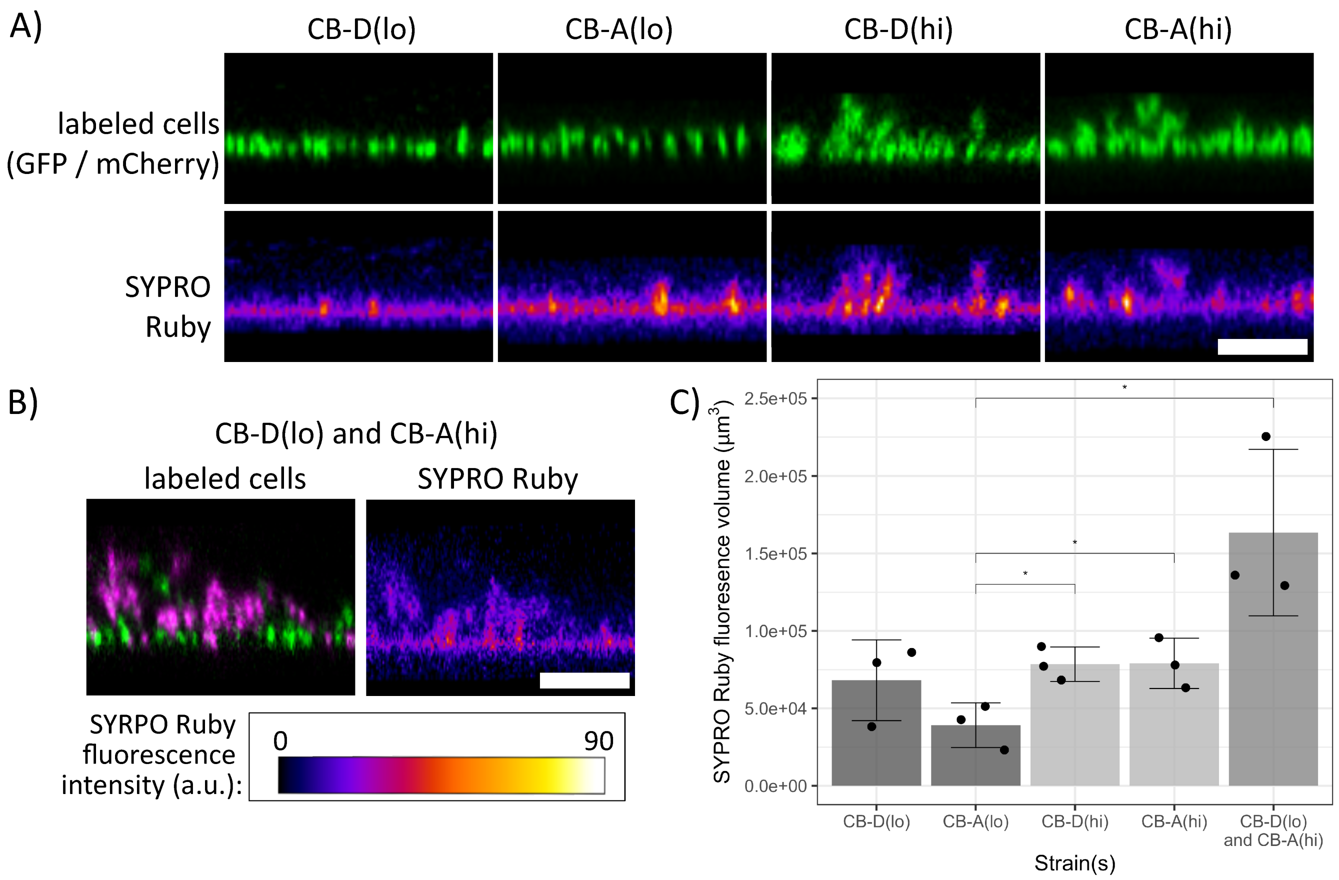

**Figure S4. Biofilm matrix protein staining with SYPRO Ruby. A)** Side views of *S. pontiacus* monoculture biofilms grown for 24 h, displaying GFP signal from labeled cells (top row) and SYPRO Ruby staining (bottom row). *S. pontiacus* strains are constitutively labeled with GFP (CB-D(lo) and CB-D(hi)) or mCherry (CB-A(lo) and CB-A(hi)). **B)** Side views of mixed-induction co-culture biofilm grown for 24 h (left) and stained with SYPRO Ruby (right). *S. pontiacus* cells are constitutively labeled with GFP (CB-D(lo); green) and mCherry (CB-A(hi), magenta). Scale bars (A and B): 10 μm. **(C)** Integrated volume of SYPRO Ruby fluorescence within monoculture and co-culture biofilms normalized by volume of constitutive cell fluorescence. Points represent volumes of three independent Z-stack images and boxes depict the median (bold line), 25th and 75th percentiles (box), and 1.5 times the interquartile ranges (whiskers). Student's t-tests were performed to determine statistically significant differences between means (* $p \leq 0.05$; ** $p \leq 0.01$; *** $p \leq 0.001$; **** $p \leq 0.0001$.

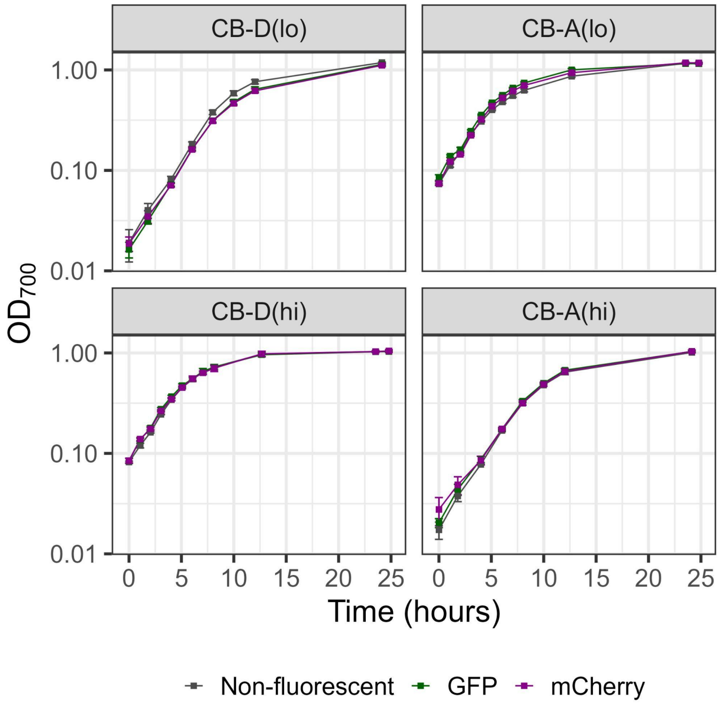

**Figure S5. Growth dynamics of constitutively fluorescent strains.** Strains were grown in SMM at 25°C with 200 rpm shaking. Points denote the mean of biological triplicates and error bars indicate standard deviation from the mean at each time point.

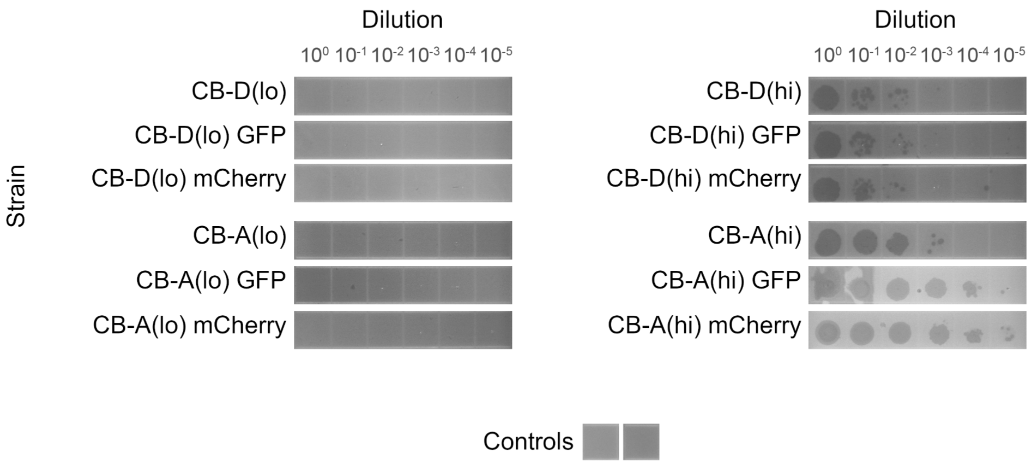

**Figure S6. Phage titers of constitutively fluorescent strains in the absence of exogenous prophage induction.** Plaque assays of phage dilutions depict titers of strains grown in SMM at 25°C overnight. Phage dilutions (10 µl) were inoculated onto host organisms susceptible to lysis by the respective phage types (CB-A(hi) for φ-D-harboring strains and CB-D(lo) for φ-A-harboring strains). Controls represent phage-free SMM medium inoculated onto hosts (left, CB-A(hi) host; right, CB-D(lo) host).

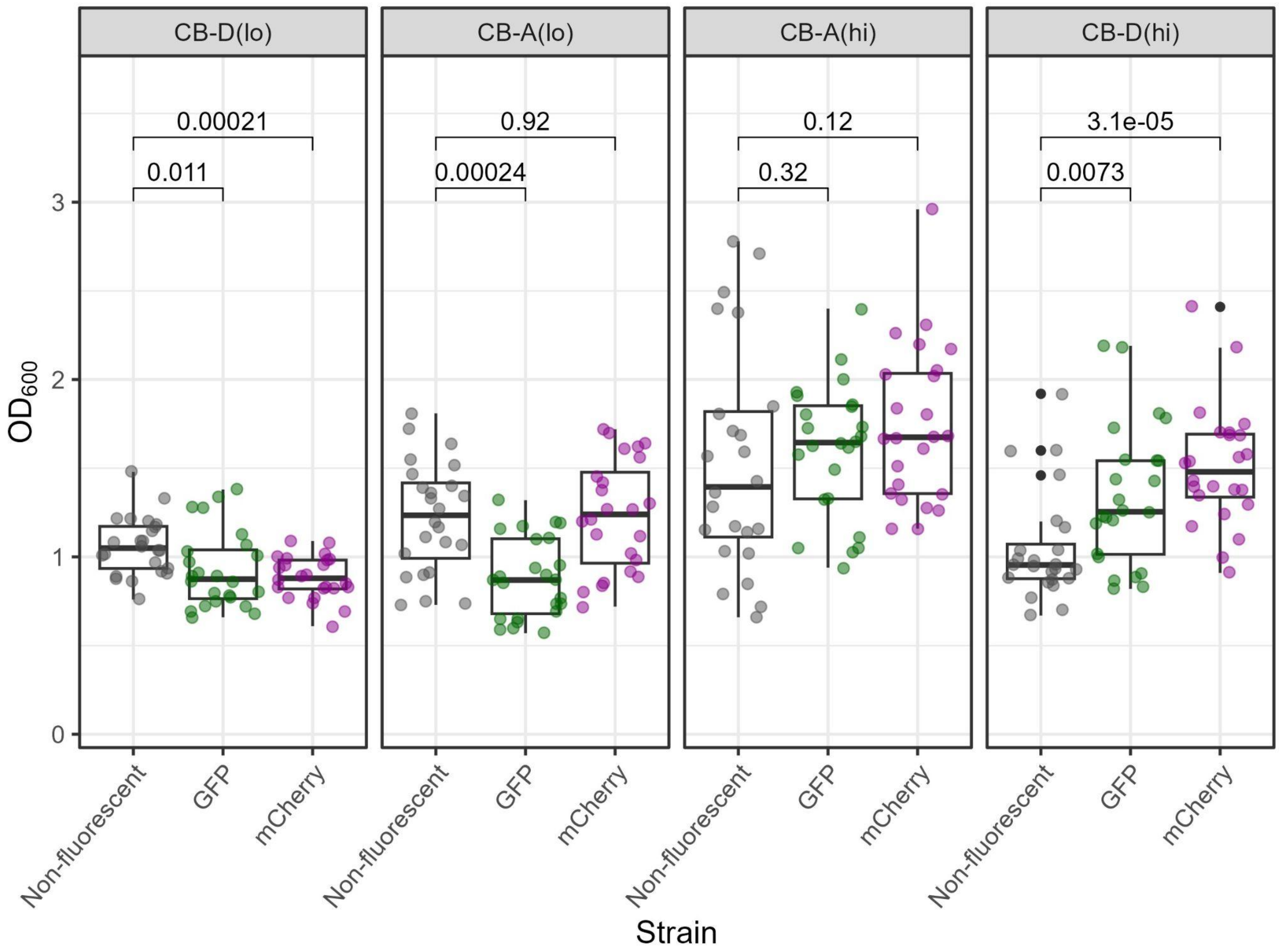

**Figure S7. Relative biofilm formation of constitutively fluorescent strains.** Crystal violet biofilm assays of strains and their fluorescent derivatives. Plots depict the median (bold line), 25th and 75th percentiles (box), 1.5 times the interquartile ranges (whiskers), and outliers (black dots) with all replicates overlaid (transparent circles of a unique color for each strain). On each panel, pairwise Wilcoxon tests were performed using the non-fluorescent strain as a reference group and p-values for each comparison are shown.
